# Epidermal Cell Dynamics Regulates Rice Lamina Joint Morphogenesis and Leaf Angle Formation through *OsZHD1* and *OsZHD2* Regulation

**DOI:** 10.1101/2025.01.20.633825

**Authors:** Yiru Xu, Heng Zhou, Xiaojiang Wu, Wuyu Cui, Shouling Xu, Xi He, Dan Xiang, Ming Zhou, Lilan Hong

## Abstract

The lamina joint is a critical determinant of leaf angle and crop architecture. While epidermal cells play a fundamental role in organ morphogenesis, influencing the overall shape and function of plants, their impact on lamina joint morphology has been largely overlooked. Here, we tracked the cellular dynamics of the rice lamina joint epidermis during leaf angle formation. We found that the asymmetric elongation between the lateral and medial edges, determined by spatial differences in the longitudinal elongation and number of epidermal cells, was a key factor in leaf angle formation. Mutations in the homeobox genes *OsZHD1* and *OsZHD2* disrupted the growth patterns of epidermal cells in the lamina joint, resulting in a decreased leaf angle. Restoring *OsZHD1* expression in the epidermis reverted the reduced leaf angle phenotype of *oszhd1 oszhd2*, highlighting the importance of epidermis development for lamina joint morphogenesis. Transcriptome data suggested that *OsZHD1* and *OsZHD2* were involved in auxin synthesis, which modulates leaf angle by inhibiting the growth of lamina joint epidermal cells. This study underscores the significance of epidermal cells in shaping the lamina joint and elucidates the critical role of *OsZHD1* and *OsZHD2* in regulating epidermal cell behavior and leaf angle formation.

## Introduction

Leaf erectness, which is determined by the leaf angle, is a critical trait for optimizing plant architecture in cereals (Qian et al., 2016; Wang et al., 2024). A smaller leaf angle facilitates efficient light interception for photosynthesis and improves canopy aeration, ultimately contributing to increased yield under high-density planting conditions (Sakamoto et al., 2006; Cao et al., 2022; Liu et al., 2022). In rice (*Oryza sativa*), the leaf angle is primarily determined by the lamina joint, which connects the leaf blade and the leaf sheath (Zhou et al., 2017). The morphology of the lamina joint is governed by its cytological structures, which are regulated by both hormonal signals and environmental factors (Zhou et al., 2017; Xu et al., 2021; Cao et al., 2022).

The well-ordered cellular organization provides the basic structure of the lamina joint. Transverse section analyses reveal that the internal tissue of the rice lamina joint primarily consists of parenchyma cells, sclerenchyma cells, vascular bundles, and aerenchyma. Parenchyma cells form the basic structural framework, while sclerenchyma tissues offer essential mechanical support. The leaf angle results from a balance between the pushing force from the expanding parenchymal cells at the adaxial side of the lamina joint and the supporting force from the structural components of the lamina joint. The vascular bundles and aerenchyma facilitate the transport of water and nutrients while ensuring air circulation (Zhou et al., 2017; Wang et al., 2020; Cao et al., 2022).

Based on the morphological features and cytological behaviors of the lamina joint, its development can be divided into six successive stages (Zhou et al., 2017; Liu et al., 2024). Systemic observations of the lamina joint at different developmental stages reveal distinct cytological changes. Global transcriptome analyses indicate that these cytological changes are accompanied by corresponding shifts in gene expression at each stage of lamina joint development (Zhou et al., 2017; Wang et al., 2020). During the early stages, active cell proliferation is supported by the expression of genes related to cell cycle regulation. In subsequent stages, the formation of different tissues such as aerenchyma and vascular bundles is driven by the upregulation of genes involved in cell wall biosynthesis and differentiation.

Altered leaf inclination can result from abnormal cell composition, disrupted cell division, or irregular cell elongation (Sun et al., 2015; Huang et al., 2021; Huang et al., 2023). Hormonal signals and environmental factors are crucial for regulating leaf angle by altering the cytological structures of the lamina joint (Luo et al., 2016; Xu et al., 2021; Cao et al., 2022). Several plant hormones, including brassinosteroids (BRs), auxins, gibberellins, and cytokinins, are involved in this regulation (Sun et al., 2015; Yano et al., 2019; Huang et al., 2023). Environmental changes can trigger hormone-mediated responses that modulate cell growth at various stages of lamina joint development (Xu et al., 2021). Among these regulators, auxin biosynthesis and signaling are critical for controlling leaf inclination. Auxin has a negative regulatory role in leaf angle formation, as reduced auxin levels lead to enlarged leaf angles by promoting parenchyma cell division and elongation at the adaxial side (Zhao et al., 2013; Yoshikawa et al., 2014; Zhang et al., 2015). Treatment of the rice flag leaf lamina joint with 1-naphthaleneacetic acid (NAA, a synthetic auxin) can reduce flag leaf angle (Huang et al., 2021). Auxin also regulates the secondary cell wall biosynthesis of sclerenchyma cells; reduced cell wall thickness in these cells leads to an exaggerated leaf angle, as they fail to adequately support the leaf, resulting in bending away from the vertical axis (Huang et al., 2021).

Although previous studies have demonstrated the involvement of multiple factors in lamina joint development and systematically revealed how the growth dynamics of different tissues in the lamina joint determine leaf angle, there remains a notable lack of cytological observations on the epidermal cells of the lamina joint. The development of the epidermis is critically important for controlling organ morphology. As the outermost tissue, the epidermis provides structural integrity; its rigidity and arrangement significantly contribute to the overall architecture of the plant, helping to maintain its shape and stability (Javelle et al., 2010; Zuch et al., 2022). Furthermore, epidermal cells can either limit or drive the growth of internal tissue cells within organs (Savaldi-Goldstein et al., 2007). The epidermis also plays a key role in regulating organ growth through various plant hormones, such as BRs and ethylene, serving as the primary site for their action (Vaseva et al., 2018; Kelly-Bellow et al., 2023).

Here we concentrated our observations on the epidermal morphology of the lamina joint. The epidermis of the mature lamina joint resembles an irregular quadrilateral when viewed from the side (facing the leaf angle). To investigate the lamina joint morphogenesis in detail, we established a nomenclature for its four edges: the edge closest to the new leaf is the lateral edge, while the edge farthest from the new leaf is the medial edge; the blade edge is at the junction of the lamina joint and the leaf blade, and the sheath edge is at the junction of the lamina joint and the leaf sheath (Figure 1A). We hypothesize that epidermal development plays a critical role in leaf angle formation and we utilized live imaging and image processing techniques to track the growth and division behavior of rice lamina joint epidermal cells. Based on this hypothesis, we predict that abnormalities in the development of the lamina joint epidermis will influence the size of the leaf angle.

**Fig.1.**
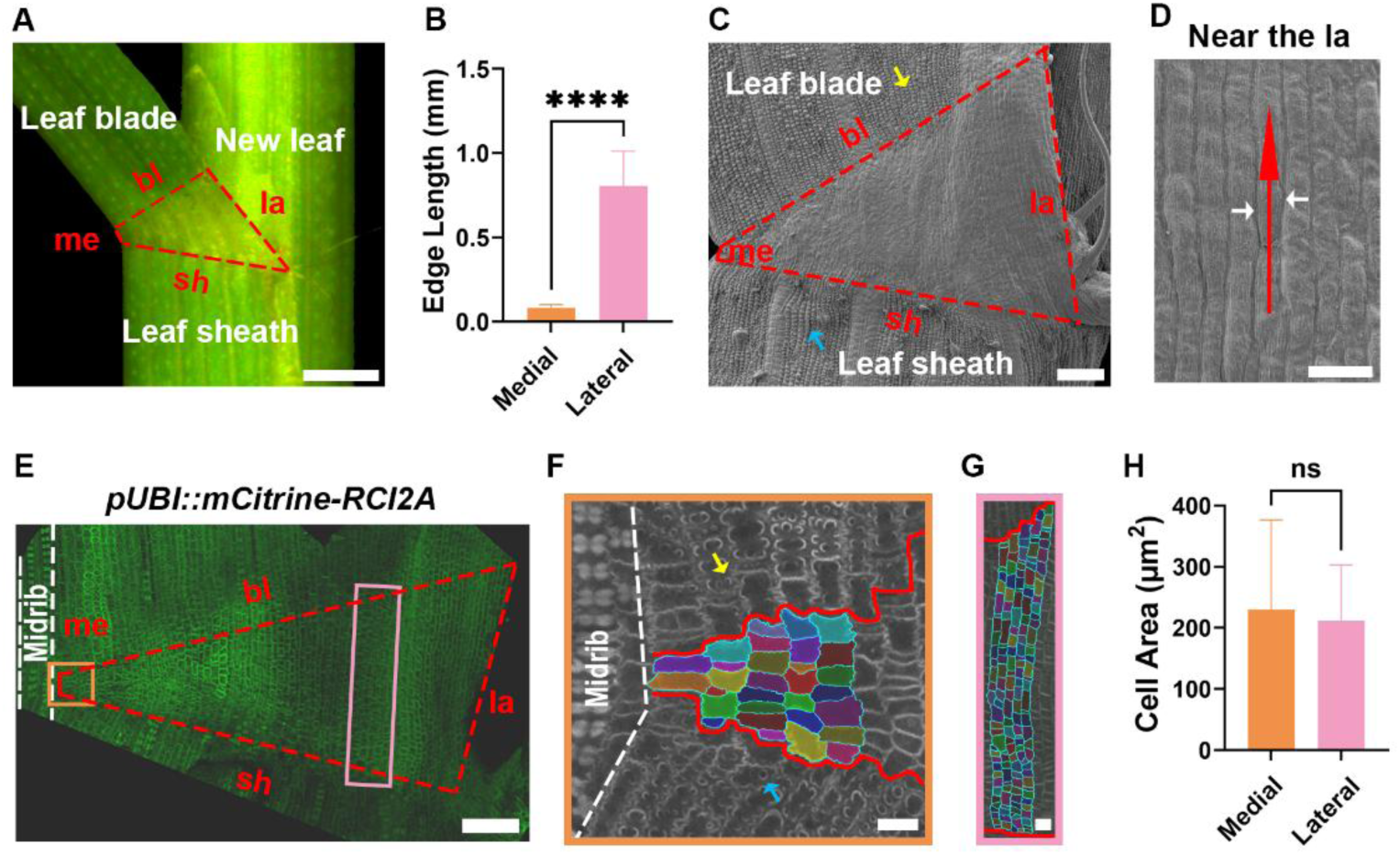
The lateral edge of the rice lamina joint contains more epidermal cells, resulting in a greater length compared to the medial edge. **(A)** Morphological diagram of the lamina joint epidermis. The side view surface of the lamina joint consists of four edges: the lateral edge (la, closest to the new leaf), and the medial edge (me, farthest from the new leaf), the sheath edge (sh, located at the junction between the lamina joint and the leaf sheath) and the blade edge (bl, located at the junction between the lamina joint and the leaf blade). Scale bar, 0.5 mm. **(B)** Lengths of the lateral edge and the medial edge of the lamina joint from the second complete leaf of 4-week-old seedlings. Data are presented as mean ± SD (n = 7). Statistical significance is determined using Student’s t-test: ****P < 0.0001. **(C**, **D)** Scanning electron micrographs of the lamina joint epidermis. **(C)** shows the entire epidermis of the lamina joint under scanning electron microscopy. The red dashed lines outline the lamina joint area. The yellow and blue arrowheads indicate silicon deposits attached to the leaf blade and leaf sheath, respectively. Scale bar, 100 μm. **(D)** shows cells near the lateral edge, with the red arrow pointing to the cell files, while the white arrows indicate borders between cell files. Scale bar, 25 μm. **(E**-**G)** Confocal images of the lamina joint using a plasma membrane reporter (*pUBI::RCI2A-mCitrine*). **(E)** shows the entire lamina joint captured by laser confocal microscopy. The white dashed lines highlight the midrib region adjacent to the medial edge. Scale bar, 100 μm. **(F)** is a magnified view of the orange box in **(E)**. The region between the solid red lines is the lamina joint. There are only a few cells in each file on the medial edge of the lamina joint. Yellow and blue arrowheads denote silicon deposits on the leaf blade and leaf sheath, respectively. Scale bar, 20 μm. **(G)** is a magnified view of the pink box in **(E)**, also with the lamina joint outlined by solid red lines. The number of cells per file near the lateral edge is considerably greater than that at the medial edge. Scale bar, 20 μm. **(H)** Area comparison of epidermal cells between the medial and lateral edges shows no significant difference. Data are presented as mean ± SD (n = 70 cells for the medial edge, n = 86 cells for the lateral edge, from 3 lamina joints). Statistical analysis is performed using Student’s t-test (ns = no significance).

## Results

### The lateral edges have more epidermal cells than the medial edges in rice lamina joint

Organ morphogenesis in plants is largely confined by epidermal cells (Savaldi-Goldstein et al., 2007; Javelle et al., 2010; Vaseva et al., 2018; Zuch et al., 2022; Kelly-Bellow et al., 2023). However, very little is known about the role of epidermal cells on lamina joint morphogenesis in grass. We aimed to address this issue in rice and imaged the lamina joint of the second complete leaf of 4-week-old rice seedlings using scanning electron microscopy (SEM), in order to obtain an overview on the morphology of epidermal cells in the rice lamina joint (Figure 1, C and D). Our observations revealed that the surface of the leaf blade and sheath epidermal cells was sprinkled with silica deposits, whereas the surface of the lamina joint epidermal cells was relatively smooth and lacked silica deposits. This difference in the abundance of silica deposits is a prominent feature distinguishing lamina joint epidermal cells from those of the leaf blade and leaf sheath (Figure 1C). Although neatly arranged cell files can be clearly seen under the SEM, it remains challenging to differentiate each cell individually (Figure 1D).

To clearly visualize the epidermal cells of the lamina joint, we created a transgenic rice line with a plasma membrane reporter (*pUBI::mCitrine-RCI2A*) and performed static imaging on dissected lamina joints of the second complete leaf from 4-week-old reporter line seedlings. Using confocal laser scanning microscopy (CLSM), we conducted z-stack scanning to capture three-dimensional (3D) images of the lamina joint. MorphoGraphx software was employed to specifically detect the fluorescence signals of the epidermal cells and segment the epidermal cells (Figure 1E). Our analysis revealed an obvious difference in the number of epidermal cells between the medial edge and lateral edge of lamina joints: the medial edge contained only a few cells (Figure 1F), while the lateral edge had dozens (Figure 1G). Given that the lateral edge is longer than the medial edge at this developmental stage (Figure 1B), and there is no significant difference (Figure 1H) in the area of lamina joint epidermal cells, we suggest that the larger number of cells along the lateral edge is a key factor contributing to this difference.

### The increased leaf angle results from the asymmetric elongation between the lateral and medial edges

Based on previous reports, the developmental processes of the rice second complete leaf lamina joint are divided into six successive stages (Zhou et al., 2017; Wang et al., 2020; Liu et al., 2024). Stage 1 to 3 are defined as the stages of organogenesis, during which the lamina joint is enveloped by the sheath of the previous leaf. Stages 4 to 6 correspond to the stages of leaf angle formation, during which the lamina joint is exposed to the air and begins to bend, forming a curvature. To elucidate the controlling mechanism of leaf angles, we focused on observing the epidermal morphology changes of lamina joints during the process of leaf angle formation, spanning from the emergence of the lamina joint (the beginning of stage 4, when the seedling is 15-day-old and the lamina joint of the second complete leaf begins to be exposed to air) to the maturity of the lamina joint (the end of stage 6, when the seedling is 27-day-old and the leaf angle of the second complete leaf reaches its maximum). The transverse size of the lamina joint reaches its maximum at stage 4 (Zhou et al., 2017). Our findings also showed that from the emergence to maturity, the lamina joint mainly changes the length of the lateral edge, suggesting that the increase in the length of the lateral edge is the primary cause of the lamina joint’s morphological changes (Figure S1).

To explore how the spatiotemporal behaviors of lamina joint epidermal cells impact the leaf angle size, we live-imaged and tracked the dynamics of lamina joint morphogenesis on the cellular level during the formation of the leaf angle using the *pUBI::mCitrine-RCI2A* seedlings. The lamina joint epidermal cells of the second complete leaf were imaged at three time points from the emergence to maturity: the 1^st^ day (1D), the 7^th^ day (7D), and the 13^th^ day (13D). On the 1^st^ day, the lamina joint was just exposed to air and began forming the leaf angle. By the 7^th^ day, the leaf angle had reached half of its mature size. By the 13^th^ day, the leaf angle had approached its maximum size (Figure 2A). Due to the technical limitations of live imaging in rice lamina joint, such as the low signal-noise ratio of fluorescent signals resulting from the strong autofluorescence in rice tissue, it was difficult to capture the growth dynamics of all the epidermal cells. We selected cells from two representative regions on the epidermis for analysis: the region between the medial and lateral edge of the lamina joint (the middle region) and the region near the lateral edge of the lamina joint (the lateral region) (Figure 2A). Cell behaviors in these two regions were analyzed using the MorphoGraphx software. During the formation of the leaf angle, the epidermal cells of the lamina joint did not undergo cell division (Figure 2B; Supplemental Figure S2A). As the leaf angle gradually increased, epidermal cells in both the middle and lateral regions exhibited a significant increase in cell area (Figure 2, C and D; Supplemental Figure S2, B and C). The epidermal cells were neatly arranged in files from 1D to 13D. We defined the direction along the cell files as longitudinal and the direction perpendicular to the cell files as transverse. We quantified the cell growth along the longitudinal direction and the transverse direction. The epidermal cells primarily elongated in the longitudinal direction, with almost no growth in the transverse direction (Figure 2, E and F; Supplemental Figure S2, D and E). In 13D, with the leaf angle reaching its mature size, there was no significant difference in cell areas between different regions (Figure 2G).

**Fig.2.**
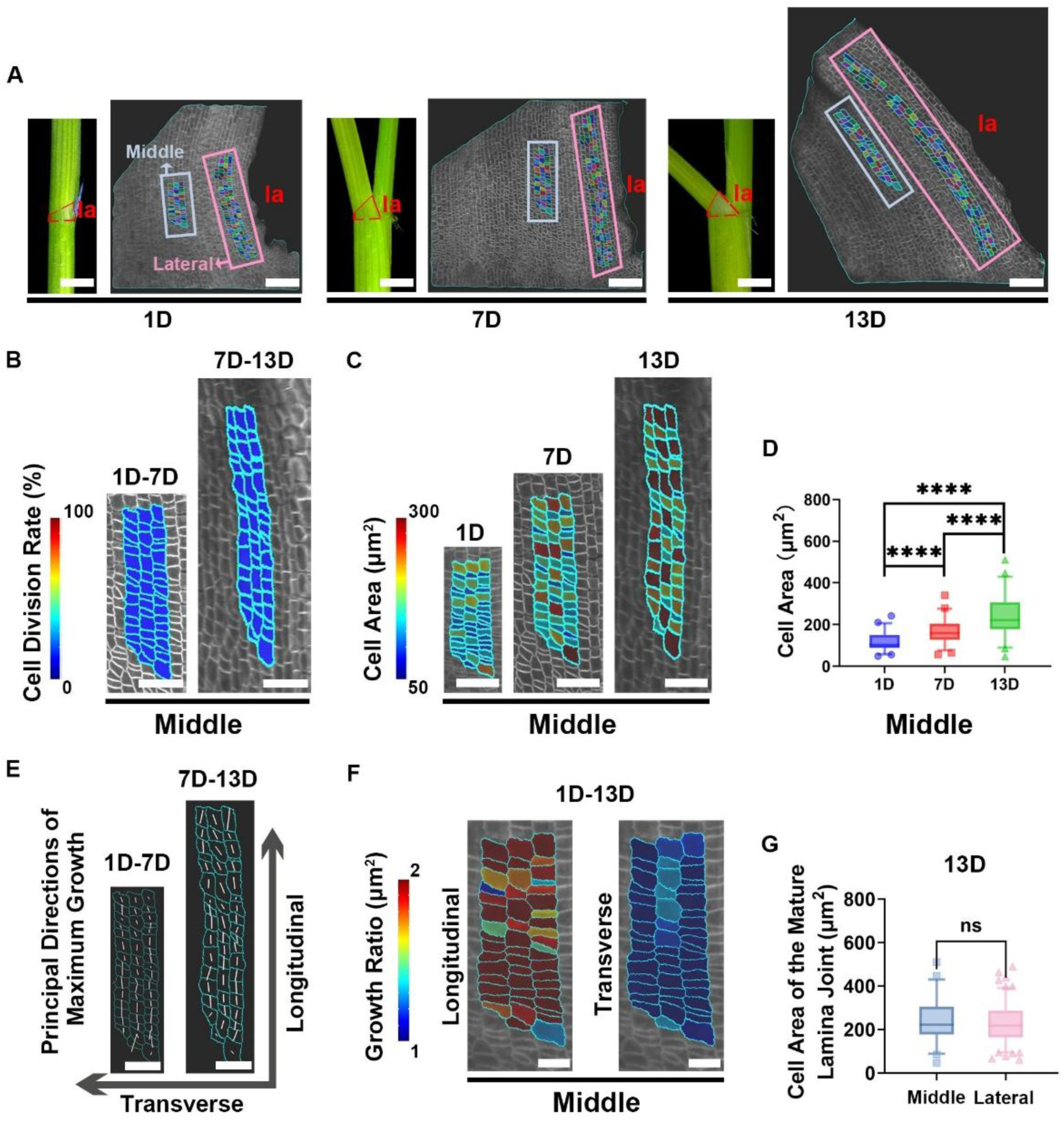
The epidermal cells of the lamina joint elongate along the cell files during leaf angle formation. **(A)** Real-time imaging of the lamina joint growth. The morphology of the lamina joint was observed on the 1D, 7D, 13D, and the epidermal cells were visualized under a confocal laser scanning microscope at these corresponding time points. The lamina joint epidermis was segmented into cells, and lineages were tracked using MorphoGraphX. Cells derived from the same mother cell at the starting time point were labeled with the same color. The red dashed lines outline the lamina joint in the stereoscope images on the left, while the blue and pink boxes mark the middle and lateral regions in the confocal images on the right, respectively. Scale bars: 1 mm for stereoscope images, 100 μm for confocal images. la, the lateral edge. **(B)** The cell division rate in the middle region, represented by the percentage of cells that divided during each growth interval (shown at the later time point). During the leaf angle formation, the epidermal cells of the lamina joint do not undergo division. Scale bars, 50 μm. **(C)** Heatmaps of cell area in the middle region. As the leaf angle increase, cells in the middle region exhibit a significant increase in area. Scale bars, 50 μm. **(D)** Boxplots showing the epidermal cell area as in **(C)**. Statistical analysis is conducted using Student’s t-test: ****P < 0.0001. n = 50 cells. **(E)** The principal directions of maximum growth (PDGmax) for epidermal cells are indicated by white lines (shown at the later time point). The longitudinal axis is parallel to the axis of the cell files, while the transverse axis is perpendicular to it. During leaf angle formation, the epidermal cells of the lamina joint primarily elongate along the direction of the cell files. Scale bars, 40 μm. **(F)** The area growth rate of cells in the longitudinal and transverse directions from 1D to 13D of observation (shown at 1D). Scale bars, 20 μm. **(G)** Boxplots of cell area for the middle and lateral regions. After the complete formation of the leaf angle, there is no significant difference in area between the epidermal cells in these two regions. n = 134 cells for the lateral region, n = 50 cells for the middle region. Statistical analysis is conducted using Student’s t-test (ns = no significance).

Although no significant difference was observed in the area of epidermal cells in different regions after the maturation of the lamina joint, we noted that, during the early stages of leaf angle formation, the cell area in the lateral region was smaller than that in the middle region (Figure 3A). This was consistent with the observation that the growth rate of cells in the lateral region was faster than in the middle region (Figure 3B). Based on these findings, we speculated that there are spatial differences in the elongation of epidermal cells of the lamina joint. The growth rate of epidermal cell area near the medial edge may be slower than that on the lateral edge, as suggested by the above-mentioned hypothesis. However, due to the limited scanning depth of confocal laser microscopy, it is challenging to dynamically image the epidermal cells on the medial edge. Therefore, we imaged the dissected lamina joint at both the emergence and maturity stages. As anticipated, we found that during the emergence stage, the cell area on the medial edge of the lamina joint was larger than that on the lateral edge, and the cell area on the medial edge did not increase as the lamina joint matured. In contrast, epidermal cells on the lateral edge had smaller areas during the early stages of the leaf angle formation, and their area significantly increased as the leaf angle increased (Figure 3, C and D). These findings highlight the spatial differences in cell elongation at different positions of the lamina joint epidermis (Figure 3E).

**Fig.3.**
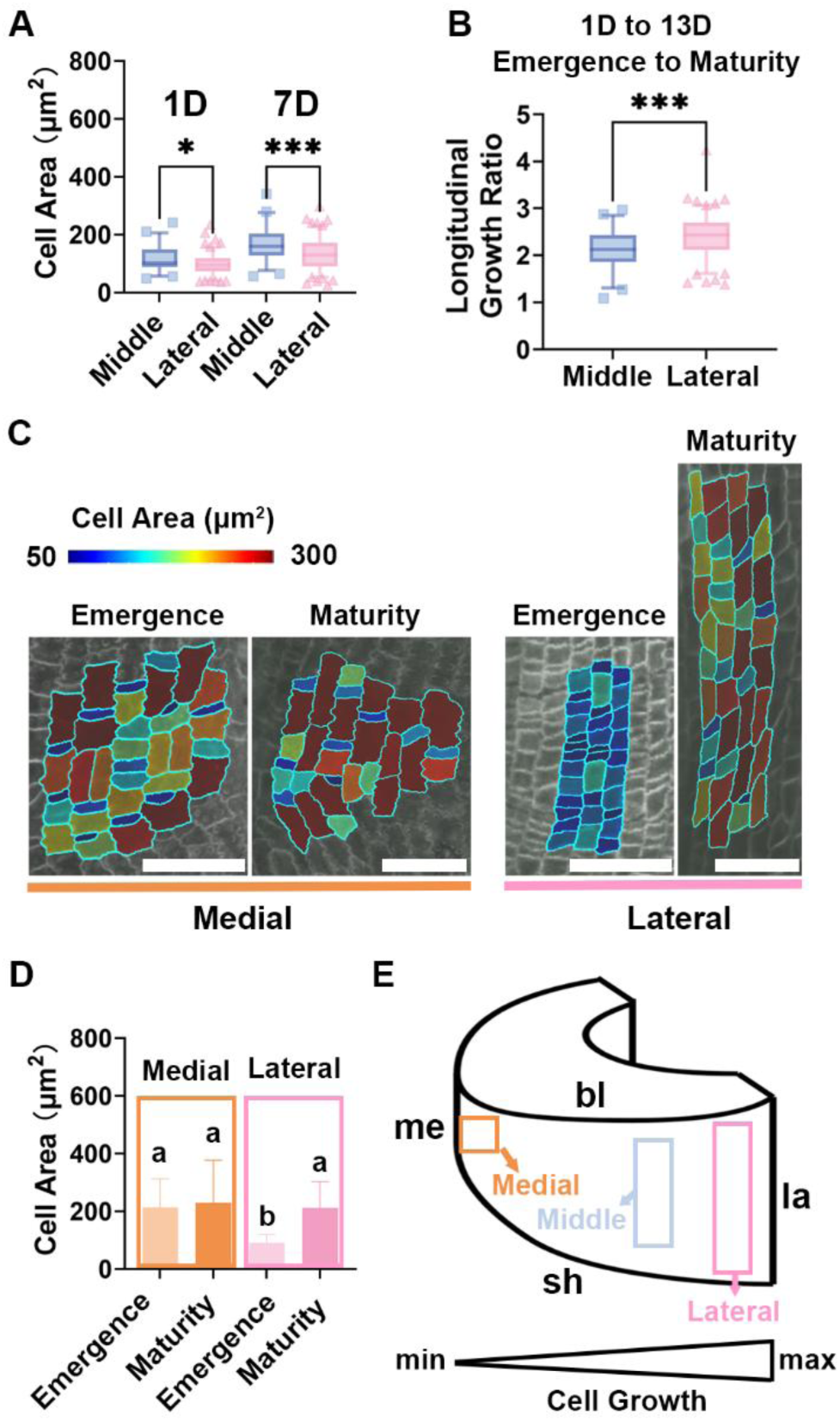
Spatial differences in the growth of epidermal cells in the lamina joint. **(A)** Boxplots of cell area for the middle and lateral regions at 1D and 7D. At both time points, the cell area in the lateral region is smaller than that in the middle region. n = 134 cells for the lateral region, n = 50 cells for the middle region. Statistical analysis is conducted using Student’s t-test (*P < 0.05, ***P < 0.001). **(B)** Boxplots of longitudinal cell growth ratio from 1D to 13D. The growth rate of cells in the lateral region is faster than in the middle region. n = 134 cells for the lateral region, n = 50 cells for the middle region. Statistical analysis is conducted using Student’s t-test (***P < 0.001). **(C)** Heatmaps showing the epidermal cell area on the medial and lateral regions at the emergence or maturity stage of the lamina joint. Scale bars, 50 μm. **(D)** Epidermal cell areas data from **(C)**. The epidermal cells on the lateral region show a significantly increased cell area during leaf angle formation while the epidermal cells on the medial edge do not elongate. At the emergence stage, epidermal cells on the lateral region are significantly smaller than those on the medial region. By the maturity stage, there are no significant difference in cell areas between the lateral and medial regions. Data are presented as mean ± SD. n = 102 cells on the medial region at the emergence stage, n = 68 cells on the medial region at the maturity stage, n = 100 cells on the lateral region at the emergence stage, n = 86 cells on the lateral region at the maturity stage. Statistical significance is determined using Student’s t-test; different letters indicate significant differences (P < 0.05). **(E)** 3D schematic diagram of the lamina joint. The orange, blue and pink boxes represent the medial, middle, and lateral regions, respectively. Spatial differences in cell elongation are observed across different regions, with the cell growth ratio gradually increasing from the medial region to the lateral region.

Considering that during leaf angle formation, the epidermal cells of the lamina joint cease division and mainly elongate along the direction of the cell file, we conclude that this longitudinal elongation results in an increase in the overall lengths of the lamina joint epidermis in the longitudinal direction. Due to the significantly larger number and greater growth ratio of epidermal cells along the lateral edge compared to the medial edge, the lateral edge undergoes more elongation than the medial edge, leading to dramatic morphological changes in the lamina joint. This asymmetric elongation between the lateral and medial edges is the key contributing factor to the formation of the leaf angle.

### Mutants lacking OsZHD1 and OsZHD2 exhibit small flag leaf angles

To investigate the molecular mechanisms regulating the patterns of lamina joint epidermal development and leaf inclination, we screened RNA-seq databases covering different stages of lamina joint development (Wang et al., 2020). Our analysis revealed that the ZF-HD family transcription factor OsZHD1 and its closest homolog OsZHD2 were potentially involved in lamina joint epidermal development. *OsZHD1* and *OsZHD2* are expressed throughout the entire stages of lamina joint development, and can significantly affect the rice architecture. (Zhou et al., 2017; Wang et al., 2020). Notably, overexpression of *OsZHD1* induces drooping leaf in rice. Both *OsZHD1* and *OsZHD2* have been demonstrated to regulate the size of bulliform cells in rice, indicating these two genes are associated with epidermis development (Xu et al., 2014). Previous research also reported that OsZHD1 and OsZHD2 regulate the cellular behavior of different organs, such as roots and flowers, by participating in different hormone pathways (Yoon et al., 2020; Yoon et al., 2022). To further investigate the role of OsZHD1 and OsZHD2 in modulating epidermal cell behaviors and leaf angle formation, we generated *oszhd1*, *oszhd2* and *oszhd1 oszhd2* knockout lines using CRISPR/Cas9 technology (Figure 4A; Supplemental table 1). We observed decreased flag leaf angles in both *oszhd1* (21.00° ± 7.18°) and *oszhd2* mutants (34.25° ± 7.66°), with a more pronounced flag leaf angle phenotype in the *oszhd1 oszhd2* double mutant lines (8.00 ± 2.99°) compared to ZH11 (71.83° ± 18.59°) (Figure 4, B and C), suggesting essential roles for *OsZHD1* and *OsZHD2* in regulating rice leaf angles. In addition, compared to the WT, mutant plants also exhibited other agronomic phenotypes such as reduced tillering, shorter plant height and smaller grain size (Figure 4C; Supplemental Figure S3).

**Fig.4.**
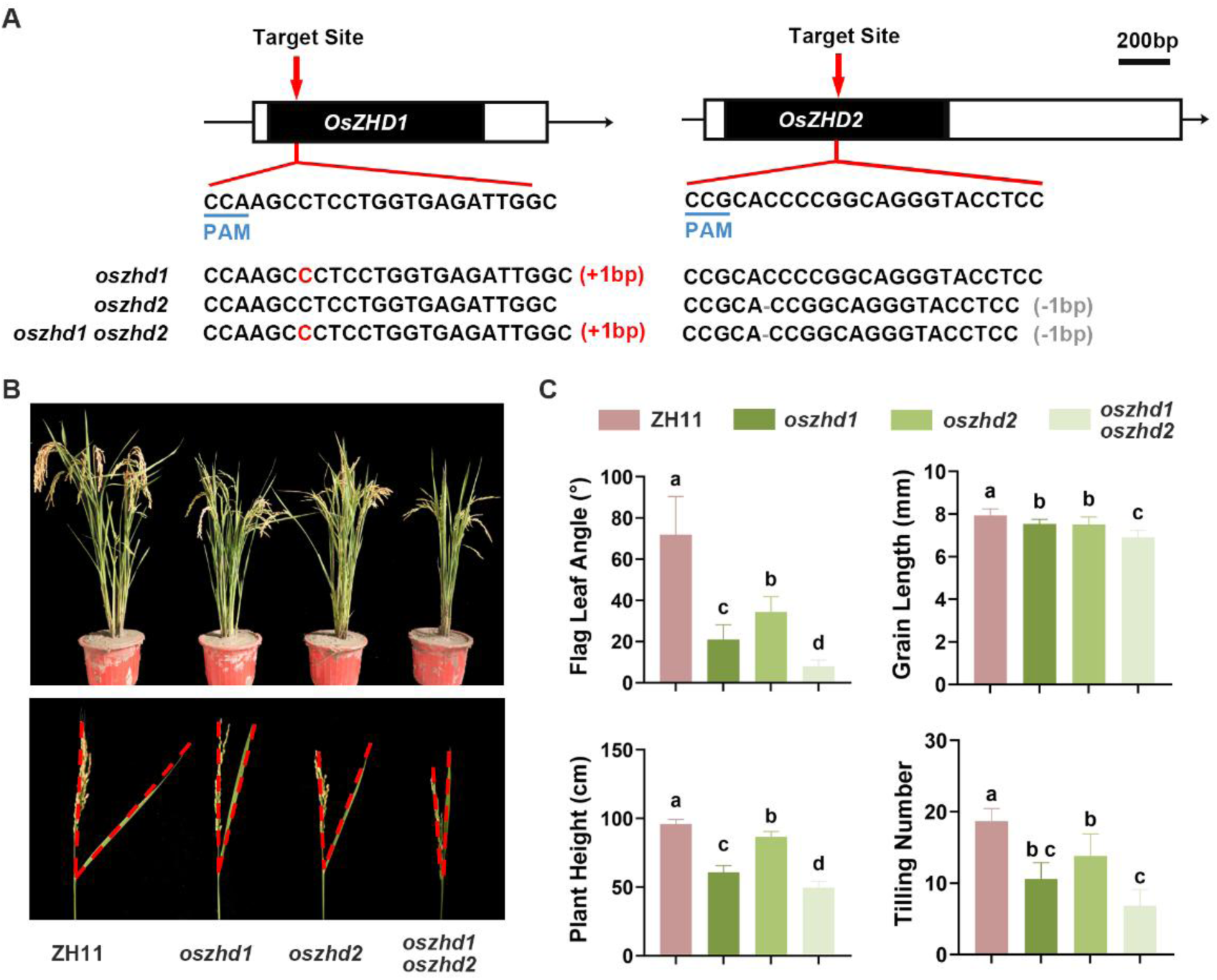
*OsZHD1* or *OsZHD2* knockout reduces leaf angle. **(A)** Target sites and mutated sequences of *OsZHD1* and *OsZHD2*. Black boxes represent exons, while arrowheads indicate the target sites of the sgRNAs. Blue underlines mark the PAM sequences. **(B)** Growth and flag leaf angle phenotypes of ZH11, *oszhd1*, *oszhd2*, and *oszhd1 oszhd2* at the ripening stage. The red dashed lines outline the angle between the flag leaves and the stems. **(C)** Statistical analysis of agronomic traits of ZH11, *oszhd1*, *oszhd2*, and *oszhd1 oszhd2*. Mutants exhibit smaller flag leaf angles, reduced tillering, shorter plant height and smaller grain size. Data are presented as mean ± SD (n ≥ 10). Statistical significance is determined using Student’s t-test; different letters indicate significant differences (P < 0.05).

### OsZHD1 and OsZHD2 affect cell proliferation and elongation in the epidermal cells of rice lamina joint

Consistent with the observed phenotypes in flag leaf angles, the mutants also exhibited smaller leaf angles at seedling stages (Figure 5, A and C). To elucidate the mechanisms by which *OsZHD1* and *OsZHD2* affect epidermal cell behavior and leaf angle formation, we analyze the morphology of the leaf lamina joints of the second complete leaves from 4-week-old seedlings (Figure 5B). We imaged them under a stereomicroscope and found that in the *oszhd1* and *oszhd2*, the total length of the lamina joint on the lateral edge was smaller than that in the WT, with *oszhd1 oszhd2* further exacerbating this difference (Figure 5D). However, no significant differences were observed in the lengths of the blade edges, sheath edges, and medial edges of the lamina joints between the different genotypes (Figure S4).

**Fig.5.**
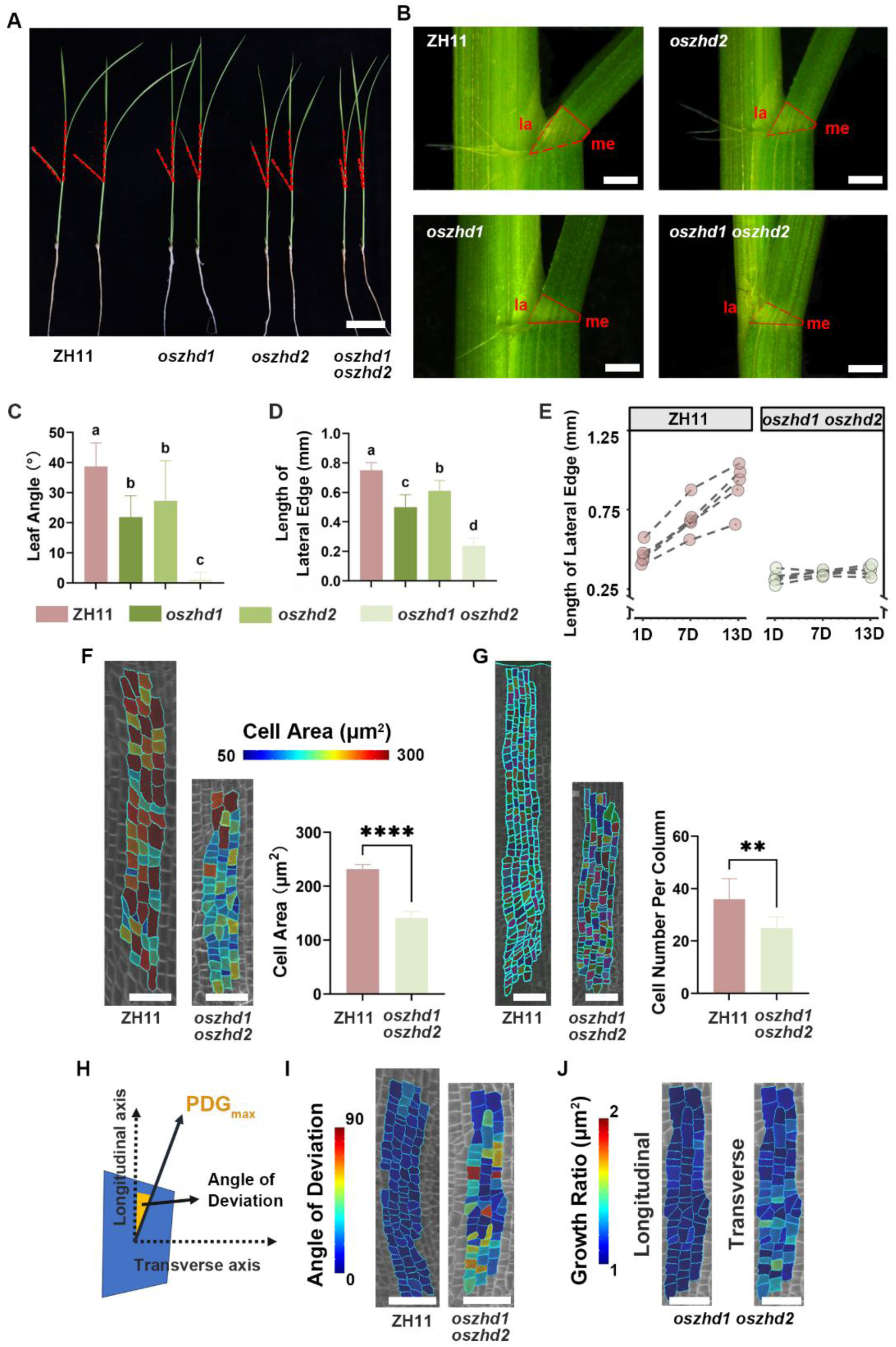
OsZHD1 and OsZHD2 affect the proliferation and elongation of epidermal cells in rice lamina joint. (A) Gross morphology of 4-week-old seedlings of ZH11, *oszhd1*, *oszhd2*, and *oszhd1 oszhd2*. The angle outlined by the red dashed lines represents the leaf angle of the second complete leaf. Scale bar, 5 cm. (B) Side views of the mature lamina joints from the second complete leaves of the seedlings shown in **(A)**. la: lateral edge, me: medial edge. Scale bars, 0.5 mm. (C) Measurement of the leaf angle of the seedlings shown in **(A)**. Data are presented as mean ± SD (n ≥ 8). Statistical significance is determined using Student’s t-test; different letters indicate significant differences (P < 0.05). (D) Measurement of the length of the lateral edge shown in **(B)**. Data are presented as mean ± SD (n ≥ 8). Statistical significance is determined using Student’s t-test; different letters indicate significant differences (P < 0.05). (E) Dynamic changes in the lateral edge length of ZH11 and *oszhd1 oszhd2*. The growth rate of the lateral edge in ZH11 is significantly faster compared to *oszhd1 oszhd2*, n = 6. (F) Heatmap of epidermal cell area of mature ZH11 and *oszhd1 oszhd2* lamina joints. The lamina joint epidermal cells in *oszhd1 oszhd2* are smaller in area compared to ZH11. Scale bars, 50 μm. Data are presented as mean ± SD (n = 505 cells for ZH11, n = 101 cells for *oszhd1 oszhd2*). Statistical analysis is performed using Student’s t-test (****P < 0.0001). (G) Epidermal cells near the fifth vascular bundle. Fewer cells are observed in the *oszhd1 oszhd2* lamina joint epidermis compared to ZH11. Scale bars, 50 μm. Data are presented as mean ± SD (n = 8). Statistical analysis is performed using Student’s t-test (**P < 0.01). (H) Schematic diagram depicting the calculation of the angle of deviation between the PDG_max_ tensor and the longitudinal axis to obtain a quantitative measure of deviation. (I) Heatmaps showing the angle of deviation of the PDG_max_ tensor from the lamina joint longitudinal axis for epidermal cells of ZH11 and *oszhd1 oszhd2*. Scale bars, 50 μm. (J) Area growth rate of cells in *oszhd1 oszhd2* in both longitudinal and transverse directions from 1D to 13D of observation (shown at 1D). Scale bars, 20 μm.

Due to the more pronounced leaf angle phenotypes in the double mutant, we focused on *oszhd1 oszhd2* for subsequent morphological analysis. We tracked the development of the lamina joints in WT and *oszhd1 oszhd2* and found that the growth rate of the lateral edge of the lamina joint in WT was significantly faster than that in *oszhd1 oszhd2* (Figure 5E). We introduced the plasma membrane reporter (*pUBI::mCitrine-RCI2A*) into *oszhd1 oszhd2*. Imaging observations of mature lamina joint epidermal cells under CLSM revealed that the area of lamina joint epidermal cells in *oszhd1 oszhd2* was smaller than that of the WT. The epidermal cell area of *oszhd1 oszhd2* is about 60% of that of the WT (Figure 5F). Additionally, fewer cells were observed in the *oszhd1 oszhd2* lamina joint epidermis. There were about 36 cells per column near the fifth vascular bundle (from the lateral edge) of the WT lamina joint, compared to only 25 cells in the counterpart region of *oszhd1 oszhd2* lamina joint (Figure 5G). While epidermal cells primarily elongated along the longitudinal direction in the WT lamina joint, the growth direction of epidermal cells in the *oszhd1 oszhd2* lamina joint was more misoriented (Figure 5H and I). In addition, the lamina joint epidermal cells in *oszhd1 oszhd2* showed low growth rates along either the longitudinal or the transverse directions (Figure 5J, compared with Figure 2F).

In summary, the mutation of *OsZHD1* and *OsZHD2* disrupts the growth patterns of epidermal cells in the lamina joint. Our results indicate that OsZHD1 and OsZHD2 may act as positive regulators of leaf angle determination by activating the proliferation and elongation of lateral epidermal cells at the lamina joint. They also regulate the direction of cell growth.

### Restoring *OsZHD1* expression in the epidermis regulates lamina joint morphology by increasing both the number and size of epidermal cells

So far, our analyses demonstrate that disrupting the growth patterns of epidermal cells affects leaf angle formation. To further test the contribution of the epidermis to leaf angle formation, we generated a transgenic rice plant expressing the full *OsZHD1* genomic sequence driven by the epidermal specific *ROC1* promoter (Ito et al., 2002; Huang et al., 2022), which restored *OsZHD1* expression in epidermal cells of the mutant. Introduction of this transgene largely rescued the reduced flag leaf angle phenotype of *oszhd1 oszhd2*, resulting in a looser plant stature compared to *oszhd1 oszhd2* (Figure 6, A, B, D, E, G). Additionally, grains of *pROC1::OsZHD1* in *oszhd1 oszhd2* plants exhibited a significant increase in length, reversing the shorter grain phenotype observed in *oszhd1 oszhd2* (Figure S5). These results indicate that *OsZHD1* can modify the morphology of both the flag leaf lamina joint and grains solely through restoring its activity in the epidermis.

**Fig.6.**
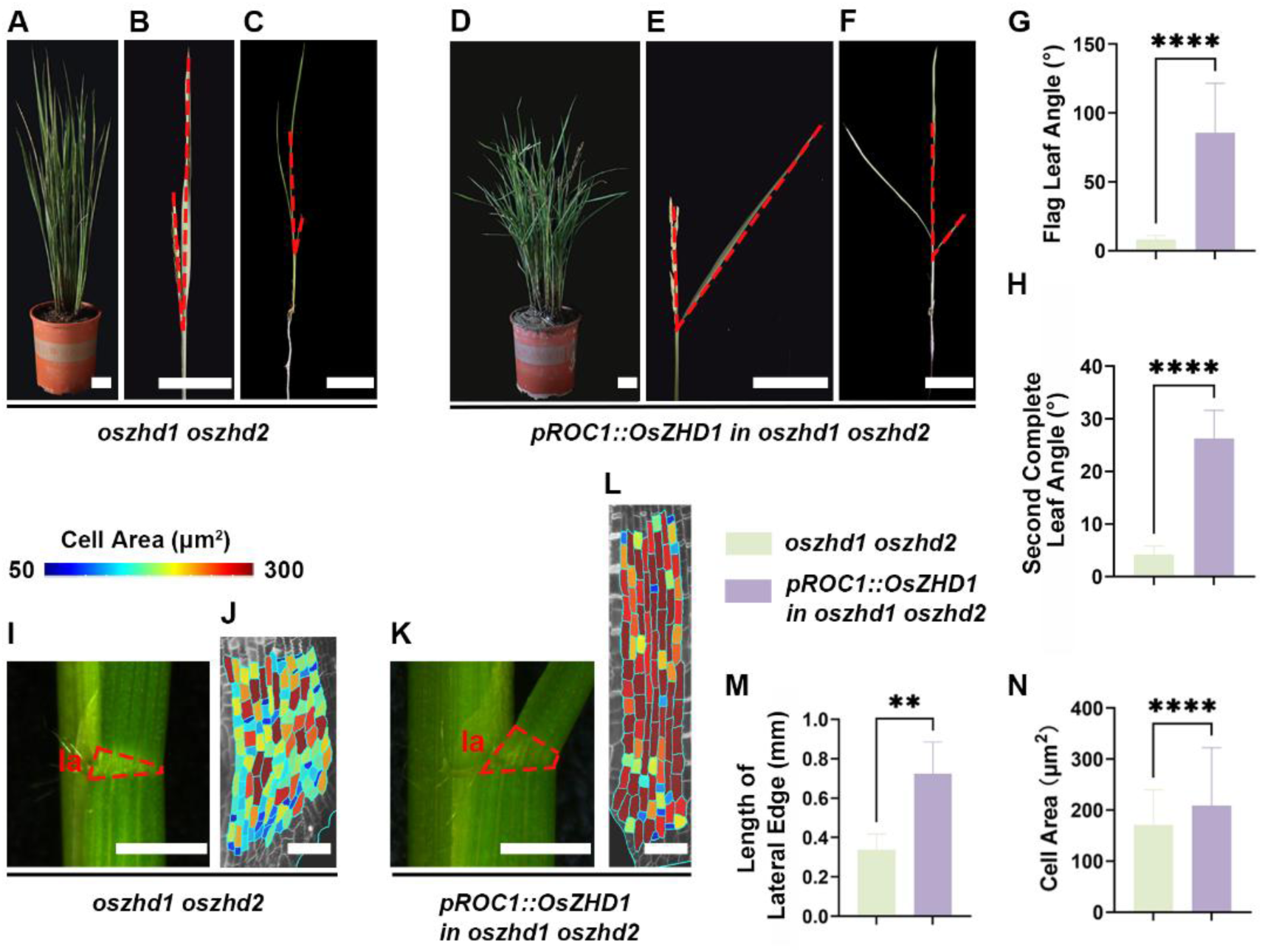
Restoring *OsZHD1* expression in the epidermis rescues the reduced leaf angle phenotype in *oszhd1 oszhd2*. **(A-C)** *oszhd1 oszhd2* exhibits a reduced leaf angle phenotype. **(A)** Growth phenotype at the ripening stage. **(B)** Flag leaf angle phenotype at the ripening stage. **(C)** Gross morphology of a 4-week-old seedling. The angle between the red dashed lines represents the leaf angle. Scale bars, 5 cm. **(D-F)** *pROC1::OsZHD1* rescues the reduced leaf angle phenotype in *oszhd1 oszhd2*. **(D)** Growth phenotype at the ripening stage. **(E)** Flag leaf angle phenotype at the ripening stage. **(F)** Gross morphology of a 4-week-old seedling. The angle between the red dashed lines represents the leaf angle. Scale bars, 5 cm. **(G)** Statistical analysis of flag leaf angles at the ripening stage. Data are presented as mean ± SD (n ≥ 10). Statistical analysis is performed using Student’s t-test (****P < 0.0001). **(H)** Measurement of the second complete leaf angles from 4-week-old seedlings. Data are presented as mean ± SD (n = 5). Statistical analysis was performed using Student’s t-test (****P < 0.0001). **(I)** Side view of the mature lamina joint from the second complete leaf of 4-week-old *oszhd1 oszhd2* seedlings. The red dashed lines outline the lamina joint. la: lateral edge. Scale bar, 1 mm. **(J)** Heatmap of the mature epidermal cell area in *oszhd1 oszhd2* as shown in **(I)**. Scale bar, 50 μm. **(K)** Side view of the mature lamina joint from the second complete leaf of 4-week-old *pROC1::OsZHD1* in *oszhd1 oszhd2* seedlings. The red dashed lines outline the lamina joint. la: lateral edge. Scale bar, 1 mm. **(L)** Heatmap of the mature epidermal cell area of *pROC1::OsZHD1* in *oszhd1 oszhd2* as shown in **(K)**. Scale bar, 50 μm. **(M)** Measurement of the length of the lateral edge shown in **(I)** and **(K)**. Data are presented as mean ± SD (n = 5). Statistical analysis is performed using Student’s t-test (**P < 0.01). **(N)** Measurement of the epidermal cell area shown in **(J)** and **(L)**. Data are presented as mean ± SD (n = 237 cells for *oszhd1 oszhd2*, n = 260 cells for *pROC1::OsZHD1* in *oszhd1 oszhd2*). Statistical analysis is performed using Student’s t-test (****P < 0.0001).

Besides the flag leaf angle, the angle of the second complete leaf from 4-week-old seedlings of *pROC1::OsZHD1* in *oszhd1 oszhd2* was also restored (Figure 6, C, F, H). Observations of the mature lamina joint morphology under a stereoscope revealed that the length of the lamina joint on the lateral edge in *pROC1::OsZHD1* in *oszhd1 oszhd2* was significantly greater compared to *oszhd1 oszhd2* (Figure 6, I, K, M). Given that *OsZHD1* acts as a positive regulator of leaf angle by promoting the proliferation and elongation of lateral epidermal cells at the lamina joint, we speculate that the number and area of epidermal cells on the lateral edge of the lamina joint in *pROC1::OsZHD1* in *oszhd1 oszhd2* were both restored, leading to an increase in the total length of the lateral edge of the lamina joint and consequently forming a larger leaf angle.

To test this hypothesis, we used Propidium Iodide (PI) staining to visualize epidermal cells in the mature lamina joint of the second complete leaf. We found that the area of epidermal cells on the lateral edge of the lamina joints of *pROC1::OsZHD1* in *oszhd1 oszhd2* was significantly increased compared to *oszhd1 oszhd2* (Figure 6, J, L, N). However, due to PI’s inability to stain the entire lamina joint, we could not accurately determine the number of epidermal cells along the lateral edge. Nonetheless, given that the total length of the lateral edge in *pROC1::OsZHD1* in *oszhd1 oszhd2* is approximately 2.14 times that of *oszhd1 oszhd2*, and that the average area of epidermal cells is roughly 1.22 times greater, we infer that the number of epidermal cells in *pROC1::OsZHD1* in *oszhd1 oszhd2* has also increased to some extent.

### OsZHD1 and OsZHD2 are involved in the regulation of leaf angle through the auxin synthesis pathway

To illuminate the underlying molecular mechanism of OsZHD1 and OsZHD2 in the regulation of lamina joint epidermal cell growth pattern and leaf angle, we generated transcriptome data of mature lamina joints from the second complete leaves for both WT and *oszhd1 oszhd2*. A total of 563 differentially expressed genes (DEGs) were identified in *oszhd1 oszhd2* compared to WT, including 299 upregulated genes and 264 downregulated genes (Figure 7A; Supplemental table 3). Kyoto Encyclopedia of Genes and Genome (KEGG) pathway analysis categorized the DEGs into diterpenoid biosynthesis, phenylpropanoid biosynthesis, cytochrome P450, limonene degradation, and tryptophan metabolism (Supplemental Figure S6). Among these, the tryptophan metabolism pathway is of particular importance due to its role in the biosynthesis of auxin, a critical plant hormone that regulates leaf angle (Cohen and Strader, 2024). The KEGG enrichment analysis results suggest that *OsZHD1* and *OsZHD2* may influence leaf angle by participating in the auxin pathway. In tryptophan-dependent pathway for auxin biosynthesis, the TRYPTOPHAN AMINOTRANSFERASE OF ARABIDOPSIS (TAA) family of tryptophan aminotransferases first converts tryptophan to indole-3-pyruvate (IPyA), while the YUCCA family of flavin monooxygenases catalyzes the conversion of IPyA to indole-3-acetic acid (IAA) (Won et al., 2011; Zhao, 2018; Matthes et al., 2019). Accumulation of auxin negatively regulates leaf angle opening (Wang et al., 2016; Huang et al., 2021). We found that the expression levels of *OsYUCCA5* and *OsYUCCA6*, two genes encoding enzymes for the rate-limiting step in auxin biosynthesis, were significantly upregulated in RNA-seq data (Figure 7B). qRT-PCR further confirmed the differential expression of these two genes between WT and *oszhd1 oszhd2* (Figure 7C). To further confirm that the auxin content was altered in *oszhd1 oszhd2*, we measured the contents of IAA and auxin precursors in the lamina joints of the second complete leaves of both WT and *oszhd1 oszhd2*. The results showed that IAA content was significantly higher in *oszhd1 oszhd2* than in WT, while IPyA content was decreased (Figure 7D). In summary, we speculate that in *oszhd1 oszhd2*, the upregulation of *OsYUCCA5* and *OsYUCCA6* promotes the conversion of IPyA to IAA. Consequently, the IPyA content decreases while the IAA content increases in the *oszhd1 oszhd2*.

**Fig.7.**
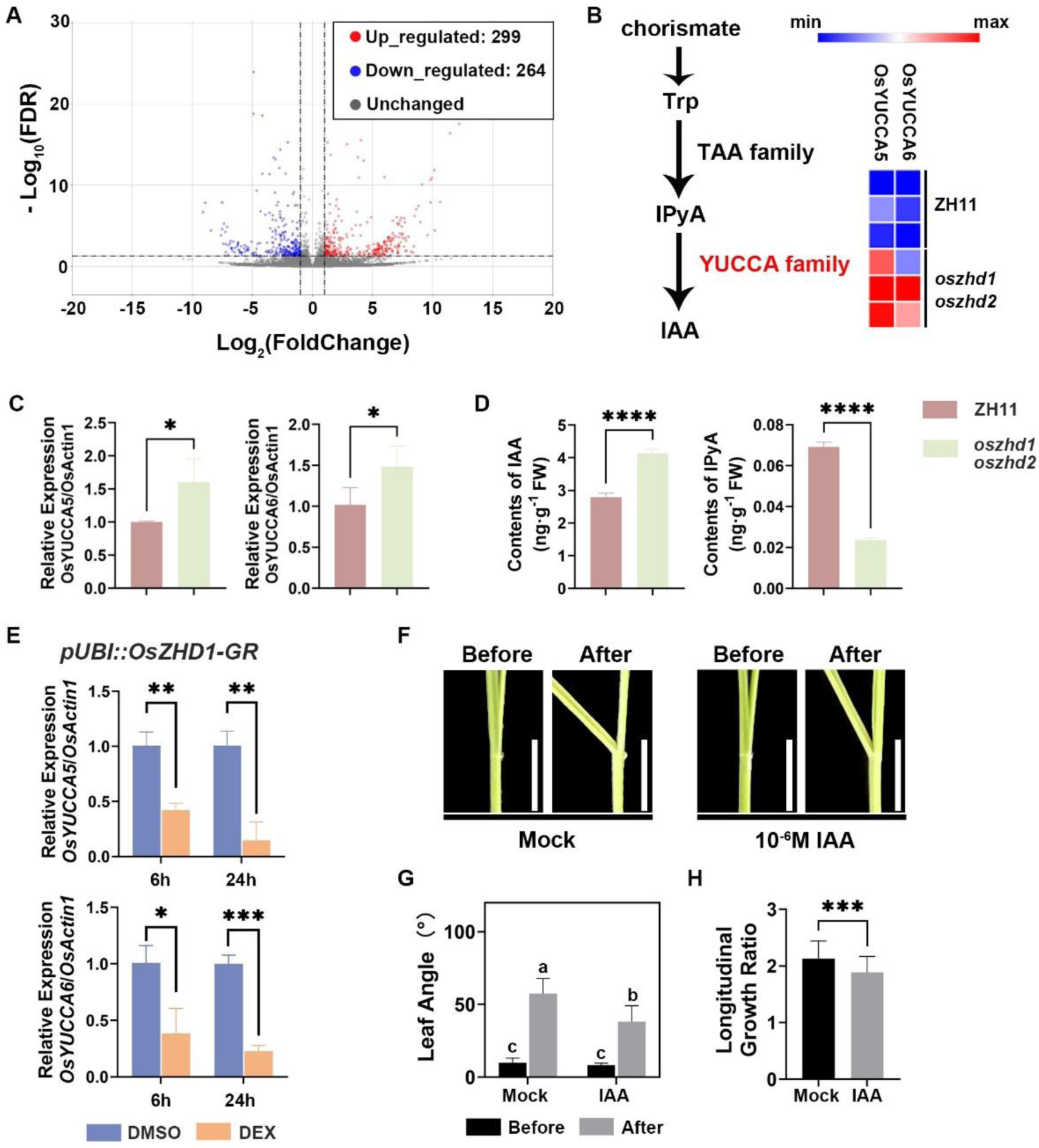
OsZHD1 and OsZHD2 regulate the expression levels of *OsYUCCA5* and *OsYUCCA6* to mediate auxin synthesis in the lamina joint. **(A)** Volcano plot of differentially expressed genes (DEGs) in the mature lamina joints of 4-week-old seedlings from *oszhd1 oszhd2* compared with ZH11. **(B)** Tryptophan-dependent pathway for auxin biosynthesis. The YUCCA family of flavin monooxygenases catalyzes the conversion of IPyA to indole-3-acetic acid (IAA). RNA-seq analysis shows increased expression levels of *OsYUCCA5* and *OsYUCCA6*, two genes encoding enzymes for the rate-limiting step in auxin biosynthesis, in the mature lamina joint of the second complete leaf of *oszhd1 oszhd2*. **(C)** Transcript levels of *OsYUCCA5* and *OsYUCCA6* analyzed by qRT-PCR. *OsActin1* served as an internal control for normalization. Data are presented as mean ± SD (n = 3). Statistical analysis is performed using Student’s t-test (*P < 0.05). **(D)** Levels of IAA and IPyA in the mature lamina joints of the second complete leaves from 4-week-old seedlings. Data are presented as mean ± SD (n = 3). Statistical analysis is performed using Student’s t-test (****P < 0.0001). **(E)** Expression levels of *OsYUCCA5* and *OsYUCCA6* in *pUBI::OsZHD1-GR* after DMSO (mock) or DEX treatment as determined by qRT-PCR. Data are presented as means ± SD (n = 3). Statistical analysis is performed using Student’s t-test (*P < 0.05, **P < 0.01, ***P < 0.001). (F) Lamina joint bending response to 10^−6^M IAA. Scale bars, 0.5 mm. (G) Quantification of the lamina joint bending assay described in (F). Data are presented as mean ± SD (n = 5). Statistical significance is determined using Student’s t-test; different letters indicate significant differences (P < 0.05). (H) Quantification of the epidermal cell longitudinal growth ratio. Data are presented as mean ± SD (n = 41 cells in the mock group, n = 50 cells in the IAA treatment group). Statistical analysis is performed using Student’s t-test (***P < 0.001).

To further verify that *OsYUCCA5* and *OsYUCCA6* are regulated downstream by *OsZHD1* and *OsZHD2*, we generated transgenic plants in which a constitutively expressed Maize Ubiquitin promoter drove the expression of a fusion gene between *OsZHD1* and the rat glucocorticoid receptor (GR) coding region (*pUBI::OsZHD1-GR*). Under dexamethasone (DEX) induction, *pUBI::OsZHD1-GR* rapidly overexpresses *OsZHD1* in the nucleus, inducing transient changes in the expression levels of downstream genes. The mature lamina joints of 4-week-old *pUBI::OsZHD1-GR* seedlings were harvested after 6 and 24 hours of DEX or DMSO treatment, and used for transcription analysis. Compared to the mock treatment, the expression levels of *OsYUCCA5* and *OsYUCCA6* were significantly downregulated after both 6 and 24 hours of DEX treatment, with levels decreasing further over time (Figure 7E). These results indicate that *OsYUCCA5* and *OsYUCCA6* are located downstream of OsZHD1. Based on our speculation that OsZHD1 and OsZHD2 have redundant functions in regulating leaf angle, we conclude that these two OsZHDs regulate the expression levels of *OsYUCCA5* and *OsYUCCA6* to mediate the synthesis of auxin in the lamina joint, thereby influencing the leaf angle phenotype.

We next tested whether auxin was sufficient to disrupt the patterns of the lamina joint epidermal development. The lamina joint of the second complete leaf from WT treated with 1 μM IAA exhibited a reduced leaf angle compared to the mock treatment (Figure 7, F and G). Following auxin treatment, the growth rate of the lamina joint epidermal cells decreased, demonstrating the negative regulatory effect of auxin on epidermal cell elongation and leaf angle formation (Figure 7H). However, the limited inhibitory effect of auxin on leaf angle suggests that *OsZHD1* and *OsZHD2* may also regulate leaf angle through other physiological processes besides inhibiting auxin biosynthesis. Previous studies have shown that BRs profoundly affect the development of the lamina joint (Sun et al., 2015; Guo et al., 2021; Liu et al., 2024). We examined the leaf bending response of the lamina joint in WT and *oszhd1 oszhd2* to BRs. Compared to WT, *oszhd1 oszhd2* appeared largely insensitive to exogenous BRs (Supplemental Figure S7). These results suggested that *OsZHD1* and *OsZHD2* might also play a role in the modulation of rice leaf angle in response to BRs. All in all, our analyses fit with a model that OsZHD1 and OsZHD2 promote the division of lamina joint epidermal cells during organogenesis and facilitate their elongation by participating in auxin biosynthesis and BR signaling during leaf angle formation, thereby influencing lamina joint morphology and regulating leaf angle (Figure 8).

**Fig.8.**
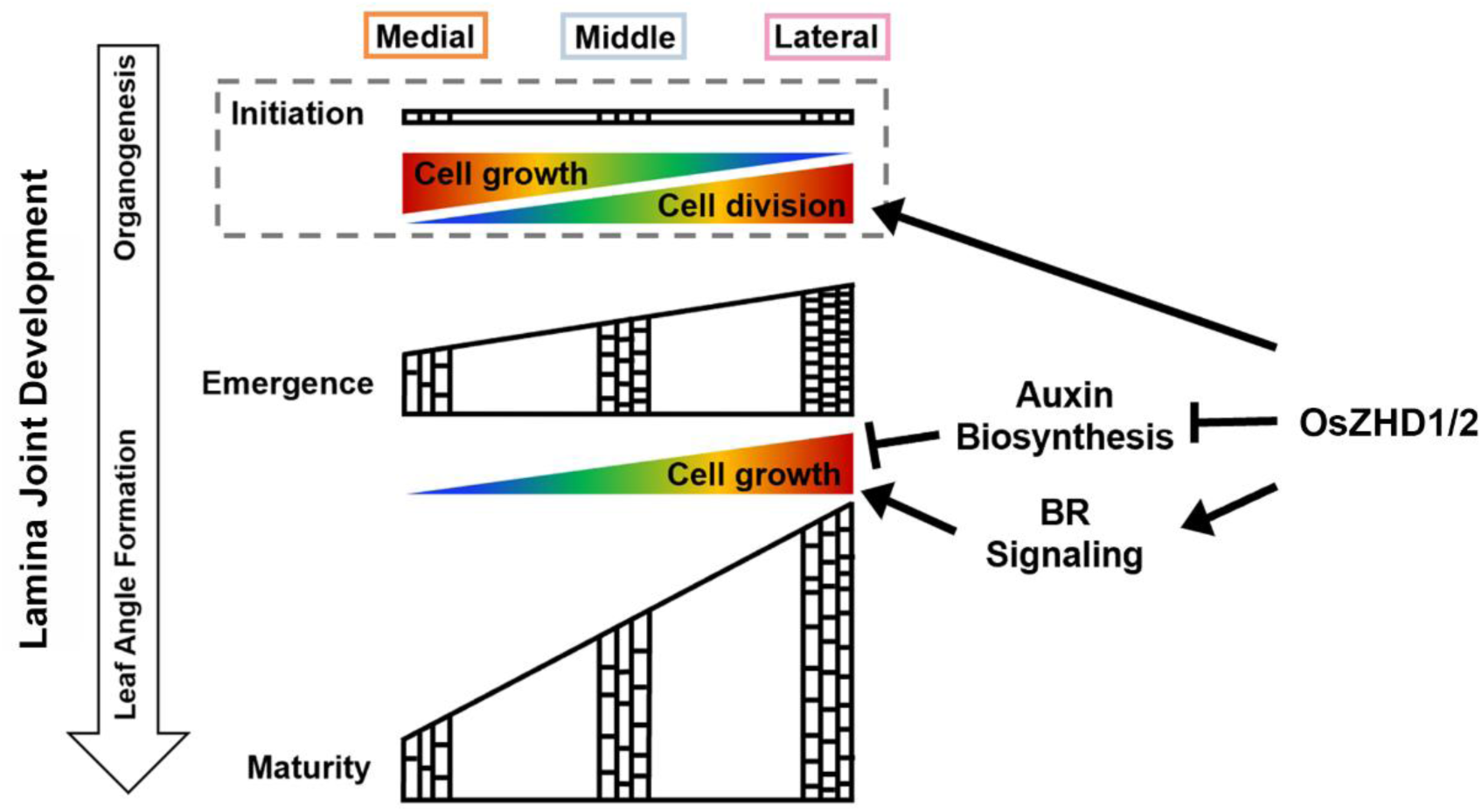
Schematic representations showing the impact of OsZHD1 and OsZHD2 on epidermal cell behavior in the rice lamina joint. This model summarizes the growth pattern of epidermal cells in the rice lamina joint and how OsZHD1/2 regulate this growth. During leaf angle formation, the epidermal cells of the lamina joint cease division and primarily elongate along the direction of the cell file. Spatial differences in cell number and elongation result in asymmetric growth between the lateral and medial edges, which contributes to leaf angle formation. Our data suggest that epidermal cell division is restricted to the organogenesis phase, during which there are spatial differences in both cell division and elongation in the lamina joint. OsZHD1/2 promote epidermal cell division during organogenesis and facilitate elongation by participating in auxin biosynthesis and BR signaling during leaf angle formation, thereby influencing lamina joint morphology and regulating leaf angle.

## Discussion

### The growth of rice lamina joint epidermal cells determines the size of the leaf angle

The lamina joint serves as an ideal model for studying the regulatory mechanisms of cytology and organ development (Zhou et al., 2017). In the study of rice lamina joint development, anatomical dissection and X-ray microcomputed tomography are commonly employed to observe the cellular morphology of internal tissues and their contribution to lamina joint morphogenesis (Wang et al., 2020; Guo et al., 2021; Huang et al., 2021; Liu et al., 2024). Despite the critical role of the epidermis in shaping organ morphology, its function and growth dynamics have often been overlooked in studies of lamina joint development. To address this gap, we generated a plasma membrane reporter (*pUBI::mCitrine-RCI2A*) to dynamically track the behaviors of lamina joint epidermal cells during leaf angle formation. Our results demonstrate that the longitudinal elongation of lateral epidermal cells contributes to leaf angle formation. Furthermore, the mature leaf angle is positively correlated with the length of the lamina joint along the lateral edge, which is determined by both epidermal cell number and cell area (Figure 1 and 2). Cytological analysis reveals that *OsZHD1* and *OsZHD2* play key roles in regulating leaf angle by promoting both cell division and elongation of epidermal cells at the lateral edge of the lamina joint (Figure 4 and 5). Notably, restoring *OsZHD1* expression in the epidermis rescues the defective leaf angle phenotype of *oszhd1 oszhd2*. This restoration is accompanied by an increase in both the number of epidermal cells and their cell area compared to *oszhd1 oszhd2*, further emphasizing the significant impact of epidermal cell development on leaf angle formation (Figure 6).

Plant cells are interconnected by their cell walls, and the coordinated growth of different cell layers and tissue types is essential for formation of physiologically efficient organs (Javelle et al., 2010; Kelly-Bellow et al., 2023). Therefore, the proper establishment of organ morphology is closely tied to the coordinated growth of epidermal and internal tissues. Previous studies have shown that lamina joint development involves multiple cellular processes within internal tissue (Zhou et al., 2017; Wang et al., 2020). Results of our dynamic observations of epidermal cells in the lamina joint is consistent with the previously reported behavior of internal cells. For instance, cell division occurs in both epidermal and internal cells before stage 4. During leaf angle formation, from stage 4 to stage 6 of lamina joint development, both epidermal and parenchyma cells undergo cell elongation (Figure 2; Supplemental Figure S2). The precise regulation of these processes collectively shapes lamina joint morphology and determines leaf angle size. Due to the suitability of the lamina joint for dynamic growth tracking of both epidermal and internal tissues, it serves as an ideal organ in rice to observe the coordination of cell behaviors during morphogenesis. Further research will focus on exploring the coordination and underlying mechanisms between epidermis and internal tissues during lamina joint morphogenesis.

### Spatial and temporal control of epidermal cell growth directs the rice lamina joint morphology

Since dynamic changes in cells determine the morphology of organs, understanding organ morphogenesis requires precisely linking the spatiotemporal behavior of cells with changes in organ morphology, which is currently a hot and challenging topic in developmental biology research (Kierzkowski et al., 2019; Peng et al., 2022). In model plants such as *Arabidopsis* and *Antirrhinum*, real-time live imaging of organ development has greatly advanced the study of the mechanisms controlling morphogenesis (Hong et al., 2016; Rebocho et al., 2017). Monocotyledonous plants have distinct morphological and anatomical structures compared to dicotyledonous plants. How monocots regulate cell growth, division, and differentiation during development to maintain specific organ morphology remains to be elucidated. To address these unresolved issues, our study specifically focused on the cellular dynamics of the rice lamina joint, developing a non-destructive method by using CLSM to investigate the detailed behavior of epidermal cells during leaf angle formation. Our research identified the lamina joint as an ideal organ for tracking cell behaviors in morphogenesis in rice for several reasons: (1) compared to leaves and sheaths, the epidermal cells of the lamina joint have less wax and silica deposition, resulting in reduced interference from plant tissue autofluorescence when using fluorescent protein reporters; (2) during most of the period in which the lamina joint undergoes morphological changes, it remains exposed rather than enclosed by other organs, making it convenient for microscopic observation and manipulation; and (3) the lamina joint is relatively small, allowing the growth of the entire organ to be observed under a high-power microscope.

In plants, the precise spatial and temporal control of cell proliferation and expansion determines the differential growth that defines organ shape and size (Serra and Perrot-Rechenmann, 2020). Our results demonstrate that during leaf angle formation, the lamina joint epidermal cells cease dividing (Figure 2; Supplemental Figure S2). Previously transcriptome analysis revealed that genes involved in stage 1 to stage 3 are primarily associated with cell division, which aligns with the cytological features observed in the lamina joint inner cells (Zhou et al., 2017; Wang et al., 2020). We deduce that cell division in the epidermal cells of the lamina joint is primarily restricted to the early stages of development, indicating the stage-specific regulation of cell division. The differences in cell numbers between the lateral and medial edges in the mature lamina joint suggest varying levels of early cell division activity at different regions of the lamina joint at the organogenesis phase. Specifically, cells near the lateral edge undergo more division, while those near the medial edge divide less (Figure 8).

In addition to the spatial differences in cell division during lamina joint organogenesis, spatial differences in cell elongation during leaf angle formation are key factors of leaf angle size. During this phase, cells near the lateral edge elongate faster, while cells near the medial edge almost cease growing (Figure 3 and Figure 8). A similar spatial pattern is observed in both inner and epidermal cells during leaf angle formation. Previous studies have shown spatial differences in lamina joint inner cell elongation: during stage 5, the abaxial parenchyma cells stop growing, while the remaining parenchyma cells on the adaxial edge continue to undergo vertical elongation (Zhou et al., 2017).

Although we were unable to image the epidermal cells of the lamina joint during organogenesis due to anatomical restriction and imaging limitations, we observed that by the emergence stage, the epidermal cells on the lateral edge were significantly smaller than those on the medial edge, despite the larger number of cells on the lateral edge (Figure 1 and Figure 3). We deduce that there are also spatial variations in the elongation of lamina joint epidermal cells during organogenesis. Specifically, we propose that the epidermal cells on the medial edge compensate for reduced cell division by promoting greater cell elongation, leading to a relatively equal length between the medial and lateral edges. This balanced edge length may facilitate better wrapping by the leaf sheath during organogenesis of the lamina joint (Figure 8).

### OsZHD1 and OsZHD2 regulate leaf angle through multiple hormone pathways

Auxin has been reported to negatively regulate the leaf angle development (Song et al., 2009; Zhang et al., 2009; Zhao et al., 2013). We found that *OsZHD1* and *OsZHD2* negatively regulate the expression of the rate-limiting enzymes of auxin synthesis *OsYUCCA5* and *OsYUCCA6*, thereby reducing auxin accumulation in the lamina joint. Since reduced auxin levels lead to enlarged leaf angles by promoting parenchyma cell division and elongation at the adaxial side (Zhao et al., 2013; Yoshikawa et al., 2014; Zhang et al., 2015), we propose that the increased IAA levels in *oszhd1 oszhd2* might inhibit the cell growth in the epidermal cells.

The regulatory effects of *OsZHD1* and *OsZHD2* on auxin accumulation may vary across different organs. *OsZHD2* can promote ethylene synthesis by positively regulating *OsACS5*, which in turn induces auxin accumulation in roots and affects the activity of rice root meristematic tissues (Yoon et al., 2020). During inflorescence meristem development, *OsZHD1* and *OsZHD2* also promote flowering by positively regulating auxin synthesis (Yoon et al., 2022). Previous studies have reported that *OsZHD1* and *OsZHD2* induce auxin accumulation by positively regulating *OsYUCCA7* in the root and inflorescence meristem, which seems to contrast with our findings that *OsZHD1* and *OsZHD2* reduce auxin accumulation by negatively regulating the expression of *OsYUCCA5* and *OsYUCCA6* in the lamina joint. However, it has been reported that most ZF-HD family transcription factors only provide DNA-binding domains and lack direct transcription regulating activity. They need to interact with other proteins to activate or inhibit the transcription of downstream genes (Tan and Irish, 2006). Moreover, previous studies have not identified any member of the YUCCA family as direct downstream targets of *OsZHD1* or *OsZHD2*. Therefore, we propose that *OsZHD1* and *OsZHD2* may exert different transcriptional regulatory effects on various direct downstream genes through interactions with different proteins, thereby influencing the expression of the YUCCA family. Considering that *OsYUCCA5* and *OsYUCCA6* can be rapidly induced by DEX in *pUBI::OsZHD1-GR* lines, we speculate that *OsYUCCA5* and *OsYUCCA6* may be potential targets of OsZHD1. The relationship of OsYUCCA5 and OsYUCCA6 with OsZHD1 in the modulation of leaf angle requires further investigation.

Emerging evidence has uncovered the mechanisms regulating cell growth on the adaxial-abaxial sides of the lamina joint, which involve the interaction of BR and auxin, at different stages of rice development (Xu et al., 2021). Here we reported that *OsZHD1* and *OsZHD2* are key transcription factors involved in BR-auxin pathways. In addition to their previously reported involvement in the ethylene and auxin pathways, we found through BR application experiments that *oszhd1 oszhd2* show reduced sensitivity to BR (Figure S7), echoing previous reports that most auxin-related mutants present changed BR sensitivity (Song et al., 2009; Zhao et al., 2013). In summary, *OsZHD1* and *OsZHD2* regulate organ development by participating in multiple hormone pathways including auxin, BR and ethylene.

## Methods and materials

### Plant materials and growth conditions

*Oryza sativa* L. ssp. *japonica* cv. ZH11 was used as the WT control for mutants throughout. *oszhd1 oszhd2* were generated using CRISPR-Cas9 technology. *oszhd1* and *oszhd2* were isolated from the progeny of an *oszhd1 oszhd2* x ZH11 cross. We obtained multiple strains with various editing types, which are listed in Supplemental Table 1. Crossing *oszhd1 oszhd2* with *pUBI::mCitrine-RCI2A* introduced the plasma membrane reporter into the *oszhd1 oszhd2* background.

Rice plants were cultivated under field conditions at experimental stations in Hangzhou (30°N, 120°E) or Changxing (31°N, 119°E) during the summer and in Lingshui (18°N, 110°E) during the winter. Seeds were soaked in distilled water at room temperature for 2 days, followed by 2 days of imbibition in water at 37℃ in a constant-temperature incubator. The germinated seeds were transferred to 96-well plates with cut bottoms and grown in water under a 16-hour light (30℃)/8-hour dark (26℃) cycle. Each plant was spaced at least 3 cm apart to allow sufficient room for the second complete leaf to fully unfold.

### Transgenic plants

To generate *pUBI::mCitrine-RCI2A*, the *mCitrine-RCI2A* sequence was amplified from *pATML1::mCitrine-RCI2A* (Hong et al., 2016) using KOD DNA polymerase (Toyobo, CAT KOD-101) with primers oSX20 and oSX21. The entire sequence was then cloned into *pCUbi1390* (Maize ubiquitin promoter inserted into *pCAMBIA1390*) digested with *KpnI* and *PstI* using T5 exonuclease-dependent assembly (Xia et al., 2019). Positive seedlings were identified using laser confocal microscopy, and strains showing strong signals were selected for subsequent experiments.

To knock out the *OsZHD1* and *OsZHD2* genes, we designed target sequences with complementary NGG PAM structures from CRISPR-P (Lei et al., 2014) for both genes and then cloned them separately into *pEntry A* (He et al., 2018) and *pEntry B* (He et al., 2018), each digested with *BsaI*. T4 DNA Ligase (NEB, CAT M0202S) was used to insert two sgRNAs into *pRHCas9* (He et al., 2018), which had been digested with *PstI* and *SpeI*. Genotyping and sequencing of the transgenic plants were performed to confirm the edit type of *OsZHD1* (using primers oYX118 and oYX125) and *OsZHD2* (using primers oYX120 and oYX126).

The 2873 bp promoter of *ROC1* was first PCR-amplified using primers oYX362 and oYX363 from WT genomic DNA, then cloned into *pRGE* (He et al., 2018), which had been digested with *PstI* and *BamHI*, to generate *pYX103*. The *OsZHD1* coding region was amplified using primers oYX386 and oYX387, and subsequently inserted into *pYX103*, digested with *SacI* and *SpeI*, to generate *pROC1::OsZHD1*.

For the construction of *pUBI::OsZHD1-GR*, the *OsZHD1* coding region was amplified using primers oYX202 and oYX190. The GR segment was cloned from *p35S::ARF3m-GR* (Xu et al., 2024) using primers oYX188 and oYX185. These two segments were fused into the *BamHI* and *SpeI* sites of the binary vector *pCUbi1390* using MultiF Seamless Assembly Mix (ABclonal, CAT RK21020).

All primers used for vector construction and genotyping are listed in Supplemental Table 2. All final constructs were verified by sequencing and transformed into the corresponding plants through *Agrobacterium*-mediated genetic transformation by Biorun Biosciences Company.

### The developmental stages of the lamina joint

The second complete leaf was selected for analysis. The developmental process of lamina joints was divided into six successive stages based on the morphological features of developing lamina joints (Zhou et al., 2017; Liu et al., 2024). The primary observations in this study focused on the emergence to maturity stages, specifically from the beginning of stage 4, when the lamina joint emerges from the previous leaf sheath and is exposed to the air, to the end of stage 6, when the leaf angle of the second complete leaf reaches its maximum.

### Scanning electron microscopy (SEM)

The samples were fixed in FAA solution (3.7% formaldehyde, 5% acetic acid, 50% ethanol), evacuated for one hour, and then preserved for more than 12 hours. The fixative was poured off, and the samples were rinsed three times with 0.1M phosphate buffer solution (pH 7.0) for 15 minutes each time. The samples were then fixed with 1% osmium acid solution for 1-2 hours. The osmium acid waste was carefully removed, and the samples were rinsed three times with 0.1M phosphate buffer solution (pH 7.0) for 15 minutes each time. The samples were dehydrated using an ethanol gradient (30%, 50%, 70%, 80%, 90%, and 95%), with each concentration applied for 15 minutes, followed by treatment with 100% ethanol for 20 minutes. Finally, the ethanol was replaced with fresh 100% ethanol. The samples were dried using a Hitachi HCP-2 critical point dryer. The dehydrated samples were coated with gold-palladium in a Hitachi Model E-1010 ion sputter for 4-5 minutes and observed under a Hitachi Model SU-8010 SEM.

### Live imaging and cell behavior analysis

To observe the development of lamina joint epidermal cells, *pUBI::mCitrine-RCI2A* was used. Under our growth conditions, it takes about 2 weeks for the second complete leaf lamina joint to reach emergence stage. At this point, the entire seedling was fixed sideways on a glass slide, and one side of the lamina joint was imaged. The imaged side of the lamina joint was marked to ensure consistent observation of the same side in future sessions. The roots of imaged samples were kept moist during the imaging to prevent the rice seedlings from drying out. Imaging was conducted using a Nikon C2si confocal microscope with a 20X objective. The imaging settings were: excitation laser at 488 nm, emission collection range of 500-550 nm, and a z-step of 0.5 μm. After imaging, the plants were returned to the growth chamber for subsequent imaging sessions. Images were processed using ImageJ (https://imagej.net/software/imagej/) to convert ND2 file to TIFF format. Growth analyses were performed with MorphographX, following previously described methods (He et al., 2020).

### RNA-seq

Seedlings were sampled by excising approximately 2 cm segments containing the mature second complete leaf lamina joint, part of leaf blade, and part of leaf sheath from both WT and *oszhd1 oszhd2*. Three biological replicates were collected, with each replicate combining lamina joints from different individual plants. Total RNA was extracted using Trizol (ABclonal, CAT RK30129) following the manufacturer’s instructions. Library preparation was conducted, and their quality was assessed using an Agilent 2100 Bioanalyzer (Agilent Technologies). Sequencing was subsequently performed on a HiSeq 2500 (Illumina) according to the manufacturer’s guidelines. Raw reads were cleaned and aligned to the ZH11 reference genome using HISAT2 (Kim et al., 2019). Genes with more than a 2-fold change in expression and a P value ≤ 0.05 were considered to be differentially expressed genes.

### Quantitative reverse transcription polymerase chain reaction (qRT-PCR)

Total RNA was extracted as described above. After removing residual gDNA, cDNA was synthesized using the PrimeScript™ RT Reagent Kit with gDNA Eraser (Takara, CAT RR047A). qPCR was performed using the Hieff^®^ qPCR SYBR Green Master Mix (Yeasen, CAT 11204ES08) on a Bio-Rad CFX96 Real-Time PCR System (Bio-Rad). Three biological replicates were conducted. *OsActin1* quantification was used as the control. All primers used for qRT-PCR are listed in Supplemental Table 2.

### DEX treatment

*pUBI::OsZHD1-GR* plants were grown in water-filled 96-well plates with cut bottoms, as described previously. It takes approximately 4 weeks for the second complete leaf lamina joint to reach stage 5. At this point, the water was replaced with either a 10 μM DEX solution or a 10 μM DMSO solution (as the mock treatment). Simultaneously, a 10 μM DEX or DMSO solution was sprayed onto the lamina joint. After 6 and 24 hours of DEX or DMSO treatment, segments of about 2 cm, containing the second complete leaf lamina joint, leaf blade, and leaf sheath, were dissected and immediately frozen in liquid nitrogen for RNA extraction and qRT-PCR analysis. The statistical analysis was conducted with Student’s t test.

### PI staining

Seedlings were sampled by excising approximately 2 cm segments containing the mature second complete leaf lamina joint, along with parts of the leaf blade and sheath, while avoiding damage to the surface of the lamina joint. The sample was placed in a 100 μg/mL PI solution for staining at room temperature for 20-30 minutes. Insufficient staining time would result in incomplete staining, while prolonged staining could cause cell death as the dye penetrates the cell membrane and binds to the nucleus. After staining, the sample was rinsed twice with sterile water to prevent background noise from excess PI dye during imaging. The rinsed sample was then placed on a slide for imaging observation.

### Quantification of phytohormones

Approximately 200 mg (fresh weight) of mature lamina joints, along with parts of the leaf blade and sheath, were collected from 4-week-old ZH11 and *oszhd1 oszhd2* plants for analysis (n ≥ 100, with 3 technical replicates). Phytohormones quantification was performed using a high performance liquid chromatography-mass spectrometry (HPLC-MS) system at Nanjing Ruiyuan Biotechnology Co., Ltd.

## Supporting information

Supplementary_Table_3_RNA-Seq_Result

## Accession numbers

Sequence from this study can be download from the rice annotation project database (https://rapdb.dna.affrc.go.jp/index.html) with the following accession numbers: *OsZHD1* (Os09g0466400); *OsZHD2* (Os08g0479400); *ROC1* (Os08g0187500); *OsYUCCA5* (Os12g0512000); *OsYUCCA6* (Os07g0437000); *OsActin1* (Os03g0718100).

## Author contributions

Conception and design of experiments, Y.X. and L.H.; experiments, Y.X., H.Z., W.C., S.X., X.H., and D.X.; live imaging and analysis, Y.X., W.C., and L.H.; RNA-seq analysis, X.W. and M.Z.; manuscript writing, Y.X. and L.H.; manuscript revising and editing, Y.X., H.Z., X.W., W.C., S.X., X.H., D.X., M.Z. and L.H.

## Acknowledgments

We thank Jingsong Bao, Dianxing Wu, Xiaoli Shu, Xiaobo Zhao and Feifei Xu for providing experimental facilities. We thank Juan Xu for providing confocal microscope. We thank Bio-ultrastructure analysis Lab of Analysis center of Zhejiang University for technical assistance in SEM experiments.

## Funding

This work was supported by the National Natural Science Foundation of China (32270867), and Hundred-Talent Program of Zhejiang University (to L.H. and M.Z.).

## Declaration of interests

The authors declare no competing interests.

## Notes

### Competing Interest Statement

The authors have declared no competing interest.

### Summary of Updates

Figure 7 revised. Supplemental files updated.

