## Supplemental_Figures_and_Legends for "Epidermal Cell Dynamics Regulates Rice Lamina Joint Morphogenesis and Leaf Angle Formation through *OsZHD1* and *OsZHD2* Regulation"

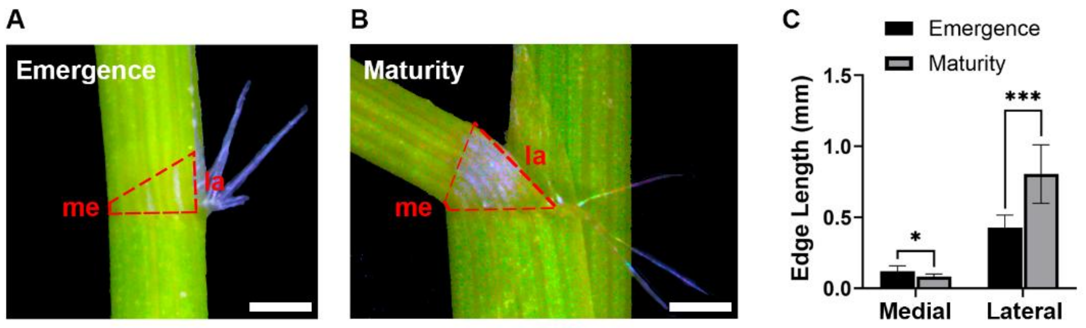


**Fig.S1 There is a significant increase in the length of the lateral edge during leaf angle formation.**

**(A)** The red dashed lines outlined the area of the lamina joint of the second complete leaf from a 15-day-old rice seedling. The lamina joint is at the emergence stage and begins to be exposed to air (the first complete leaf was removed to facilitate observation of the lamina joint in the second complete leaf). Scale bar, 0.5 mm.

**(B)** The red dashed lines outlined the area of the lamina joint of the second complete leaf from a 27-day-old rice seedling. The lamina joint is at the maturity stage, and the leaf angle of the second complete leaf reaches its maximum. Scale bar, 0.5 mm.

**(C)** Edge length analysis of the lamina joint of the second complete leaf at different developmental stages. Data are presented as mean ± SD (n = 7). Statistical significance is determined using Student's t-test: $^{*}P < 0.05$, $^{***}P < 0.001$.

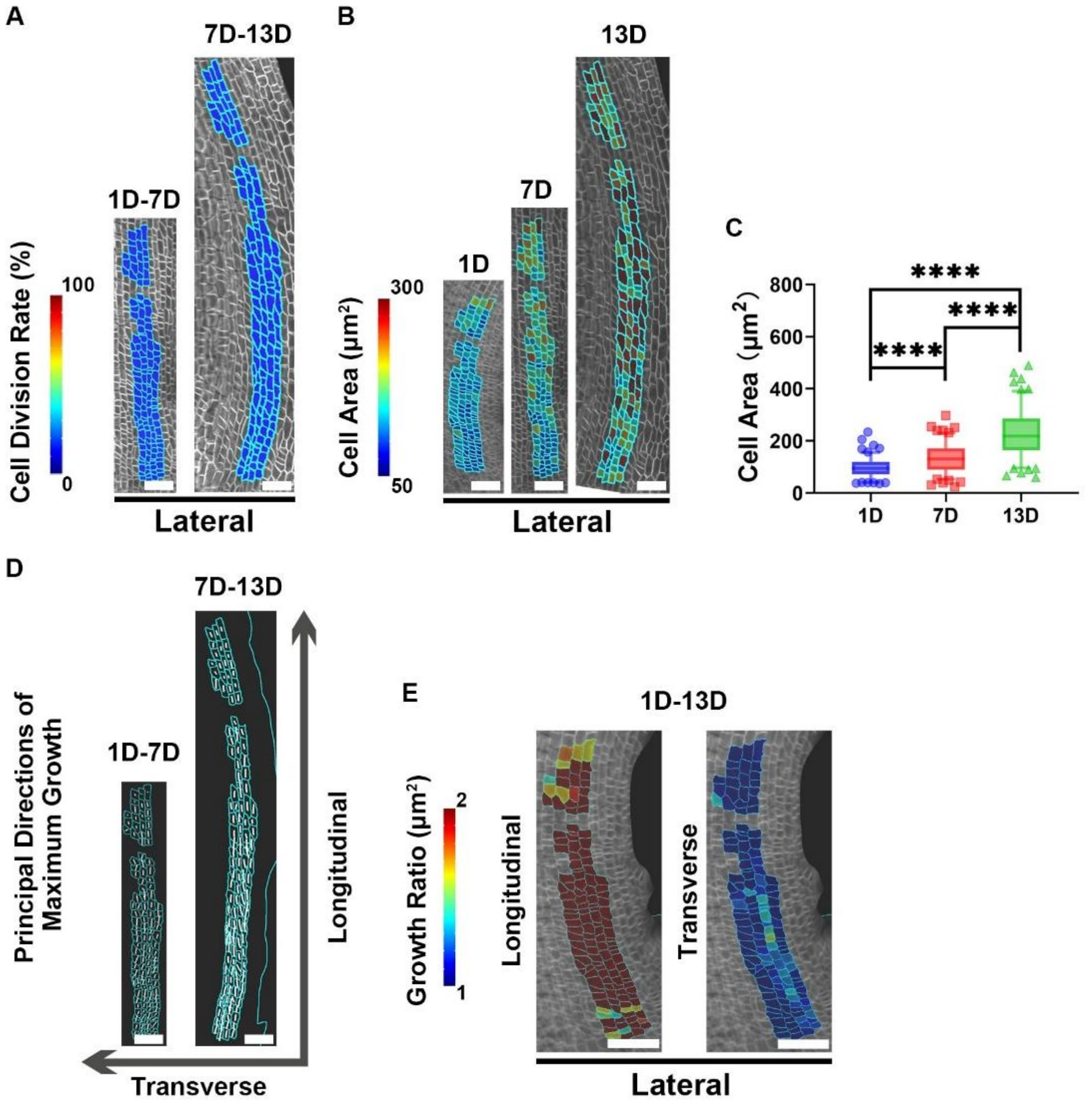


**Fig.S2 Real-time imaging of the lateral region of lamina joint epidermal cells, associated with Main Fig.2.**

**(A)** Cell division rate in the lateral region, represented as the percentage of cells dividing during each growth interval (shown at the later time point). During leaf angle formation, the epidermal cells of the lamina joint do not undergo division. Scale bars, 50 μm.

**(B)** Heatmap of cell area in the lateral region. As the leaf angle increase, the cell area in the lateral region significantly expands. Scale bars, 50 μm.

**(C)** Boxplots showing the epidermal cell area as in **(B)**. Statistical analysis is conducted using Student's t-test: $^{****}P < 0.0001$. n = 134.

**(D)** The principal directions of maximum growth (PDGmax) for epidermal cells are is indicated by white lines (shown at the later time point). During leaf angle formation, the epidermal cells of the lamina joint primarily elongate along the direction of the cell files. Scale bars, 40 μm.

**(E)** Area growth rate of cells in both longitudinal and transverse directions from 1D to 13D of observation (displayed at 1D). Scale bars, 20 μm.

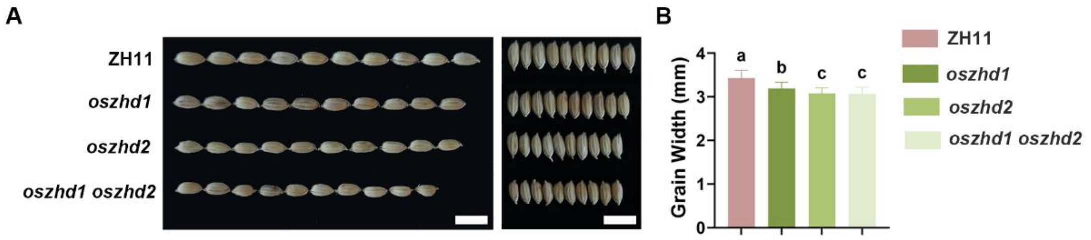


**Fig.S3 OsZHD1 and OsZHD2 knockout mutants exhibit a small grain phenotype.**

**(A)** Gross morphology of the ZH11, *oszhd1*, *oszhd2*, and *oszhd1 oszhd2* grains. Scale bars, 1 cm.

**(B)** Grain width analysis. Data are presented as mean ± SD (n = 30). Statistical significance is determined using Student's t-test; different letters indicate significant differences ($P < 0.05$).

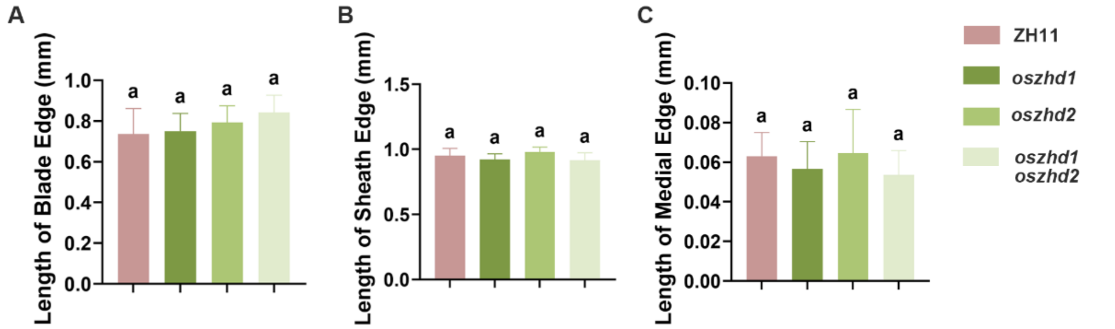


**Fig.S4 Length analysis of various knockout lines of *OsZHD1* or *OsZHD2* shown in Figure 4.**
**(A-C)** No significant differences were observed in the lengths of the blade edges **(A)**, sheath edges **(B)**, and medial edges **(C)** of the lamina joints between the different genotypes. Data are presented as mean ± SD ($n \geq 8$). Statistical significance was determined using Student's t-test; identical letters indicate no significant differences ($P > 0.05$).

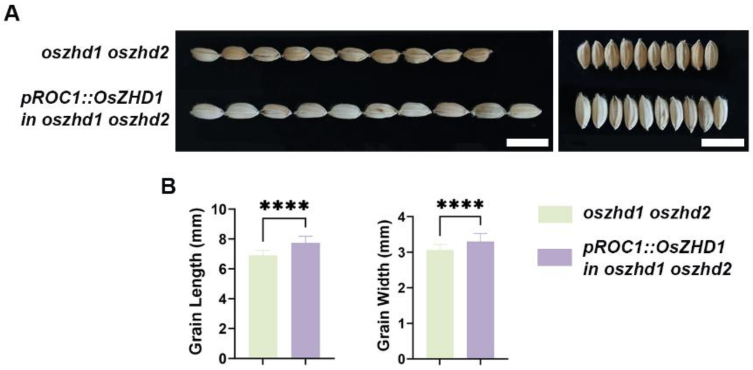


**Fig.S5 Restoring *OsZHD1* expression in the epidermis reverses the shorter grain phenotype observed in *oszhd1 oszhd2*.**

**(A)** Gross morphology of the *oszhd1 oszhd2* and *pROC1::OsZHD1* in *oszhd1 oszhd2* grains. Scale bars, 1 cm.

**(B)** Grain length and width analysis. Data are presented as mean ± SD (n = 30). Statistical analysis is conducted using Student's t-test: $^{****}P < 0.0001$.

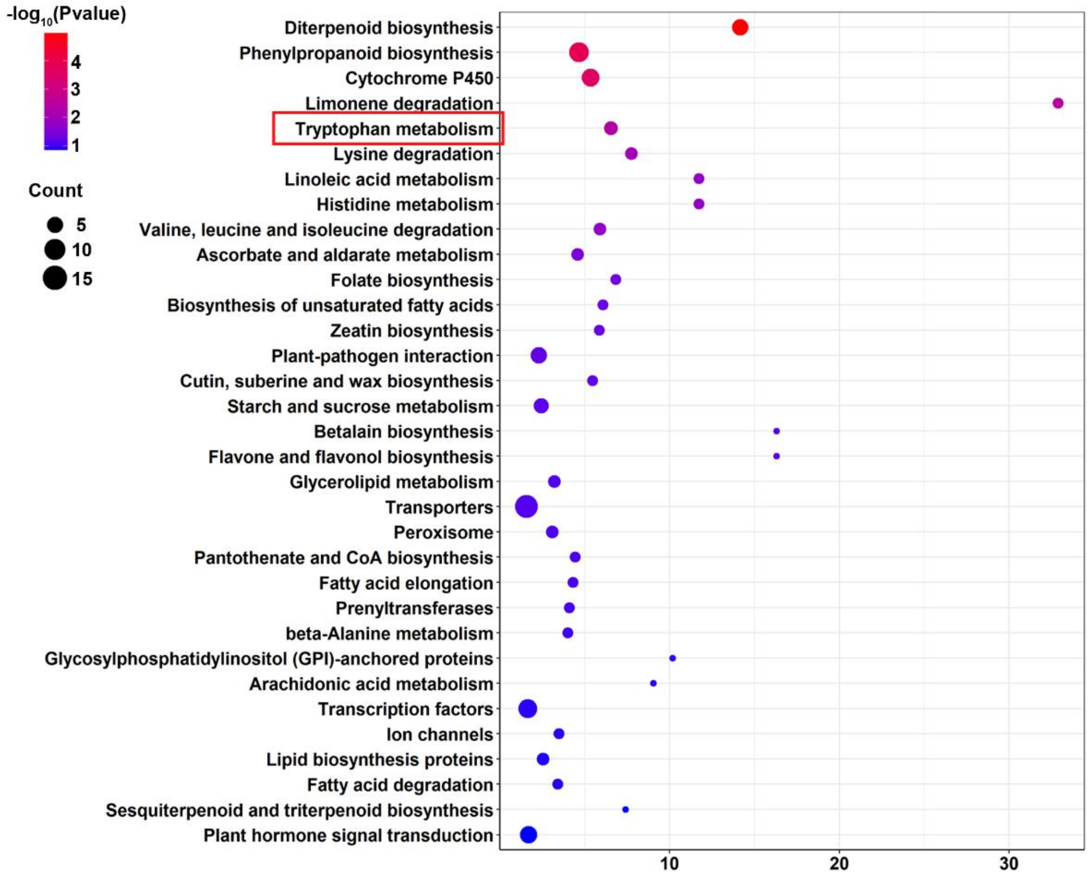


**Fig.S6 Kyoto Encyclopedia of Genes and Genome (KEGG) pathway analysis.**

KEGG pathway analysis categorized the DEGs in *oszhd1 oszhd2* into diterpenoid biosynthesis, phenylpropanoid biosynthesis, cytochrome P450, limonene degradation, and tryptophan metabolism.

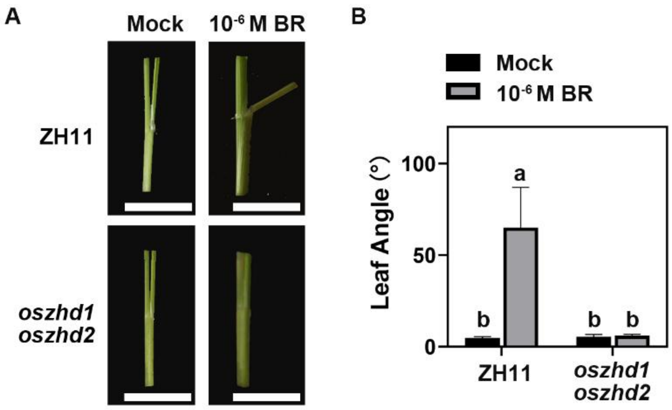


**Fig.S7** ***oszhd1 oszhd2*** **shows reduced sensitivity to BR.**

**(A)** Lamina joint bending response to $10^{-6}$M BR. Scale bars, 0.5 mm.

**(B)** Quantification of the lamina joint bending assay described in **(A)**. Data are presented as mean ± SD (n = 5). Statistical significance is determined using Student's t-test; different letters indicate significant differences ($P < 0.05$).

**Supplementary table 1. Different genotype of *oszhd1 oszhd2*.**

| | *OsZHD1* | *OsZHD2* |
|---|---|---|
| **WT** | CCAAGCCTCCTGGTGAGATTGGC | CCGCACCCCGGCAGGGTACCTCC |
| | CCAAGCCTCCTGGTGAGATTGGC | CCGCACCCCGGCAGGGTACCTCC |
| ***oszhd1 oszhd2*#1** | CCAAGC(+T)CTCCTGGTGAGATTGGC | CCGCAC(-CC)CGGCAGGGTACCTCC |
| **(+1+1/-2-2)** | CCAAGC(+A)CTCCTGGTGAGATTGGC | CCGCAC(-CC)CGGCAGGGTACCTCC |
| ***oszhd1 oszhd2*#2** | CCAAGC(-C)TCCTGGTGAGATTGGC | CCGCAC(-C)CCGGCAGGGTACCTCC |
| **(-1+1/-1-1)** | CCAAGC(+A)CTCCTGGTGAGATTGGC | CCGCAC(-C)CCGGCAGGGTACCTCC |
| ***oszhd1 oszhd2*#3** | CCAAGC(+T)CTCCTGGTGAGATTGGC | CCGCAC(-CC)CGGCAGGGTACCTCC |
| **(+1+1/-2-1)** | CCAAGC(+G)CTCCTGGTGAGATTGGC | CCGCAC(-C)CCGGCAGGGTACCTCC |
| ***oszhd1 oszhd2*#4** | CCAAGC(+T)CTCCTGGTGAGATTGGC | CCGCAC(+A)CCCGGCAGGGTACCTCC |
| **(+1+1/+1+1)** | CCAAGC(+A)CTCCTGGTGAGATTGGC | CCGCAC(+A)CCCGGCAGGGTACCTCC |

**Supplementary table 2. Primers used.**

| Primer Name | Sequence | Purpose |
|---|---|---|
| oSX20 | TTCTGCACTAGGTACCTGCAgATGGTGAG CAAGGGCGAGGAGCTGT | F_primer for amplifing mCitrine-RCI2A fragment from *pATML1::mCitrine-RCI2A* to generate *pUBI::mCitrine-RCI2A* |
| oSX21 | TTCCCGGGGATCCGTCGACCATGAAATG ATAGCGTAAGGTAT | R_primer for amplifing mCitrine-RCI2A fragment from *pATML1::mCitrine-RCI2A* to generate *pUBI::mCitrine-RCI2A* |
| oYX118 | TGCTCCTGGGTGAACTTGGT | F_primer for PCR genotyping of *OsZHD1* |
| oYX125 | GCATTTCGGCTTGACTCTT | R_primer for PCR genotyping of *OsZHD1* |
| oYX126 | TGCCACCGTAACTTCCAC | F_primer for PCR genotyping of *OsZHD2* |
| oYX120 | CGCAGACCTCCTCGCAGAA | R_primer for PCR genotyping of *OsZHD2* |
| oYX362 | gatcccccgaattactgcagAGGGTCTCTCCCTGCA TCAT | F_primer for amplifing *OsROC1* fragment from WT gDNA to generate *pYX103* |
| oYX363 | ccgagctcacccggggatccGAACGAAGCCAGGT AATTAAGC | R_primer for amplifing *OsROC1* fragment from WT gDNA to generate *pYX103* |
| oYX386 | TCggatccccgggtgagctcATGGACTTCGATGAC CATGACGA | F_primer for amplifing *OsZHD1* fragment from WT gDNA to generate *pOsROC1::OsZHD1* |
| oYX387 | cacttagcggccgcactagtTCATGGCAGCTTCTTG CCCAGGG | R_primer for amplifing *OsZHD1* fragment from WT gDNA to generate *pOsROC1::OsZHD1* |
| oYX202 | ttacttctgCACTAGGTACcATGGACTTCGATG ACCATGACGA | F_primer for amplifing *OsZHD1* fragment from WT gDNA to generate *pUBI::OsZHD1-GR* |
| oYX190 | gtttttcgagcttcGGATCCTGGCAGCTTCTTGCC CAGGG | R_primer for amplifing *OsZHD1* fragment from WT gDNA to generate *pUBI::OsZHD1-GR* |
| oYX188 | ACcTGCAGGTCGACGGATCCgaagctcgaaaaa caaagaaaa | F_primer for amplifing *GR* fragment from *p35s::ARF3-GR* to generate *pUBI::OsZHD1-GR* |
| oYX185 | TGGCTAGCGTTAACACTAGTTCATTTTTG ATGAAACAGAAGC | R_primer for amplifing *GR* fragment from *p35s::ARF3-GR* to generate *pUBI::OsZHD1-GR* |
| oYX477 | AGTGGCTCAAGGGAAGTGAC | F_primer for *OsYUCCA6* qPCR |
| oYX478 | CAGCATCTGAGGAGACACCA | R_primer for *OsYUCCA6* qPCR |
| oYX479 | attcgtgaatggctgtaggg | F_primer for *OsYUCCA5* qPCR |
| oYX480 | gtcgctctcgctgaagaact | R_primer for *OsYUCCA5* qPCR |
| OsActin1-F | CGGGAAATTGTGAGGGACAT | F_primer for *OsActin1* qPCR |
| OsActin1-R | AGGAAGGCTGGAAGAGGACC | R_primer for *OsActin1* qPCR |
